# Stress unmasks impaired endocrine coordination of glucose metabolism in dystrophin deficiency

**DOI:** 10.64898/2026.02.22.707291

**Authors:** Gretel S Major, Cara A. Timpani, Hannah Lalunio, Jiayi Chen, Lauren Boatner, Nicholas Giourmas, Stephanie Kourakis, Eva van den Berg, Ekaterina Salimova, Joel Eliades, Gang Zheng, Rong Xu, Sukhvir Kaur Bhangu, Thirimadura V. H. Mendis, Joel R Steele, Ralf B Schittenhelm, Deirdre L Merry, Louise Lantier, Christoph E. Hagemeyer, Francesca Cavalieri, Karen Caeyenberghs, Aaron P. Russell, Michael de Veer, Craig A. Goodman, Alan Hayes, Emma Rybalka, Charles P. Najt, Angus Lindsay

**Affiliations:** School of Biological Sciences, University of Canterbury, Christchurch 8041, New Zealand; Biomolecular Interaction Centre, School of Biological Sciences, University of Canterbury, Christchurch 8140, New Zealand; Institute for Health and Sport, Victoria University, Melbourne, Victoria 8001, Australia; Australian Institute for Musculoskeletal Science (AIMSS), Victoria University and Western Health, St Albans, Victoria 3021, Australia; Department of Medicine—Western Health, Melbourne Medical School, The University of Melbourne, St Albans, Victoria 3021, Australia; Department of Human Nutrition, Foods, and Exercise, Virginia Polytechnic Institute and State University, Blacksburg, VA, USA; Metabolism Core, Virginia Polytechnic Institute and State University, Blacksburg, VA, USA; Monash Biomedical Imaging, Monash University, Clayton, Victoria 3800, Australia; Monash Metabolic Phenotyping Platform, Monash Biomedicine Discovery Institute, Monash University, Melbourne, Australia; Department of Diabetes, School of Translational Medicine, Monash University, Melbourne, Victoria 3004, Australia.; Department of Chemical Engineering, The University of Melbourne, Parkville, Victoria 3000, Australia; Monash Proteomics & Metabolomics Platform, Department of Biochemistry and Molecular Biology, Biomedicine Discovery Institute, Monash University, Clayton, Victoria 3800, Australia; Department of Molecular Physiology and Biophysics, Vanderbilt University, Nashville, Tennessee, United States; Vanderbilt Mouse Metabolic Phenotyping Center, Vanderbilt University, Nashville, Tennessee, United States; School of Science, RMIT University, Melbourne, Victoria, 3000 Australia; Dipartimento di Scienze e Tecnologie Chimiche Universita’ di Roma “Tor Vergata”, Via della Ricerca Scientifica 1, Rome, 00133 Italy; Cognitive Neuroscience Unit, School of Psychology, Deakin University, Geelong, Victoria, Australia; Institute for Physical Activity and Nutrition (IPAN), School of Exercise and Nutrition Sciences, Deakin University, Geelong, Victoria, Australia; Centre for Muscle Research, Department of Anatomy and Physiology, The University of Melbourne, Parkville, Victoria 3010, Australia; Department of Medicine, University of Otago, Christchurch 8014, New Zealand; Maurice Wilkins Centre for Molecular Biodiscovery, Auckland 1010, New Zealand

## Abstract

Skeletal muscle orchestrates systemic metabolism, dynamically coordinating glucose uptake and fuel use to match energy demand. In Duchenne muscular dystrophy, loss of dystrophin is associated with altered metabolic regulation. In the mdx mouse, we show that physiological stress reveals impaired coordination between insulin and stress responses: glucocorticoid signalling increases without a proportional rise in insulin secretion, resulting in systemic hyperglycaemia despite preserved capacity for muscle glucose uptake. These data support a multi-tissue dystrophinopathy associated with altered endocrine-metabolic coordination. Skeletal muscle glycogen is elevated and incompletely mobilised under stress. The heart maintains high glucose uptake, whereas the brain exhibits reduced uptake, highlighting tissue specific differences in metabolic response. Acute insulin supplementation improves systemic glucose control and restores stress-induced behavioural deficits. Likewise, empagliflozin-mediated glucose offloading reduces stress-associated blood glucose spikes and is associated with improved muscle function to levels comparable with standard care prednisolone. These findings identify impaired coordination of endocrine and metabolic responses during stress as a contributor to metabolic vulnerability in DMD and suggest that modulating insulin availability or glucose flux can improve systemic metabolic control.

## INTRODUCTION

Skeletal muscle is a powerhouse of systemic metabolism as a driver of ∼30% of resting energy expenditure and the largest insulin-sensitive tissue in the body(*1*, *2*). The defining feature of skeletal muscle is metabolic plasticity, the ability to dynamically shift glucose uptake and fuel utilisation in response to energy availability and physical activity(*3*). This regulatory capacity is most challenged during acute physiological stress, when insulin and counter-regulatory hormones must be precisely coordinated to match tissue energy demand. Even subtle disruptions at the muscle level are sufficient to propagate widespread systemic metabolic dysfunction(*4*).

Duchenne muscular dystrophy (DMD) is an X-linked neuromuscular disease caused by mutations in the *DMD* gene(*5*). Severe and progressive muscle degeneration is characteristic of DMD, resulting in replacement of contractile tissue with fatty and fibrotic deposits. Skeletal muscle is affected first and most severely, owing to high mechanical loads and exhaustive degeneration-regeneration cycles(*6*). Cardiac muscle, with negligible regeneration capacity, undergoes progressive cardiomyocyte loss and fibrosis, resulting in fatal dilated cardiomyopathy. However, people with DMD also have high rates of systemic metabolic dysregulation including obesity, adiposity, hyperinsulinemia, insulin resistance and glucose intolerance(*7–9*). The brain and central nervous system also display characteristics of altered energy metabolism(*10–12*), predisposing people with DMD to a myriad of neurocognitive comorbidities(*13*, *14*), including a hypersensitivity to stress, coined the startle response(*15*).

Dystrophin, encoded by the *DMD* gene, is a core membrane-associated protein forming part of the dystrophin-glycoprotein complex (DGC) and anchoring the extracellular matrix and intracellular cytoskeleton(*16*). While classically associated with impaired contractile function, dystrophin deficiency is also accompanied by broad metabolic disturbances including cellular mitochondrial dysfunction(*17–21*), tissue ischemia(*22*) and hypometabolism(*23*, *24*). The transmembrane DGC acts as a key signalling hub docking receptors and enzymes that link dystrophin deficiency directly to disrupted cellular bioenergetics(*25*). For example, localisation of the insulin receptor and neuronal nitric oxide synthase to the DGC links dystrophin loss to altered insulin responsiveness(*26*) and glucose handling(*27*, *28*), respectively. Consistent with this, muscle biopsies from people with DMD show increased subcellular GLUT4 receptor abundance relative to controls(*7*), indicating dysregulated glucose utilisation rather than impaired transporter availability. Glycogen metabolism is similarly altered, with increased glycogen synthesis and storage but reduced breakdown in dystrophin-deficient skeletal muscle(*29*). Despite substantial evidence for local and systemic dysregulation of glucose and glycogen metabolism in DMD, the extent to which this reflects a primary metabolic disorder remains debated. Currently no targeted therapies address this metabolic burden. Critically, it remains unclear whether metabolic instability stems from impaired insulin availability or from peripheral insulin resistance.

Previous studies, including our own, have detailed the behavioural and physiological features of stress hypersensitivity in pre-clinical genetic models of dystrophin-deficiency(*15*, *30–36*). Here, we exploit this phenotype as a physiological stressor to unmask metabolic defects in the dystrophin-deficient *mdx* mouse model of DMD, enabling interrogation of insulin-stress coordination and metabolic flexibility in dystrophin deficiency. Using acute stress as a metabolic challenge, we observe impaired endocrine-metabolic integration, whereby increased stress signalling is not matched by a corresponding increase in insulin secretion, coinciding with reduced systemic glucose control. This occurs despite preserved insulin-stimulated glucose uptake *in vivo,* indicating that tissues retain the capacity to respond to insulin at the whole-body level and suggesting that metabolic instability is more consistent with impaired insulin availability than overt peripheral insulin resistance. Increasing insulin availability improves systemic metabolic control and alleviates stress-induced behavioural suppression, supporting a role for endocrine insufficiency in this phenotype. Empagliflozin, acting independently of insulin, diverts circulating glucose and is associated with reduced stress-induced hyperglycaemia and improved muscle function. These results indicate that metabolic vulnerability in dystrophin-deficiency is associated with altered endocrine-metabolic coordination, and that modulating insulin availability or redirecting glucose flux can improve metabolic and behavioural responses under stress.

## RESULTS

### Dystrophin deficiency is associated with heightened metabolic vulnerability to stress

To determine whether dystrophin deficiency confers heightened susceptibility to metabolic dysregulation, we investigated systemic glucose homeostasis in the absence of full-length dystrophin. Given that stress represents a potent physiological challenge to glucose homeostasis(*37*, *38*), we designed a series of experiments to investigate metabolic vulnerability. Dystrophin-deficient *mdx* mice had elevated endogenous glucose production and reduced glucose tolerance early in life (3 months), which was amplified and sustained by behavioural stress (tube-restraint intervention, hereafter stress), while wild-type (WT) mice exhibited age-dependent increases in blood glucose and reduced glucose tolerance that exceeded *mdx* levels by 12 months of age **(Fig 1A-B; Fig S1A-E)**. Independent of age, stress delayed the peak hyperglycaemic response in *mdx* mice **(Fig S1C)**. This pattern indicates an early dysregulation of both endogenous glucose regulation and glucose tolerance in dystrophin deficiency, with age-related metabolic impairment emerging later in WT mice.

**Fig. 1.**
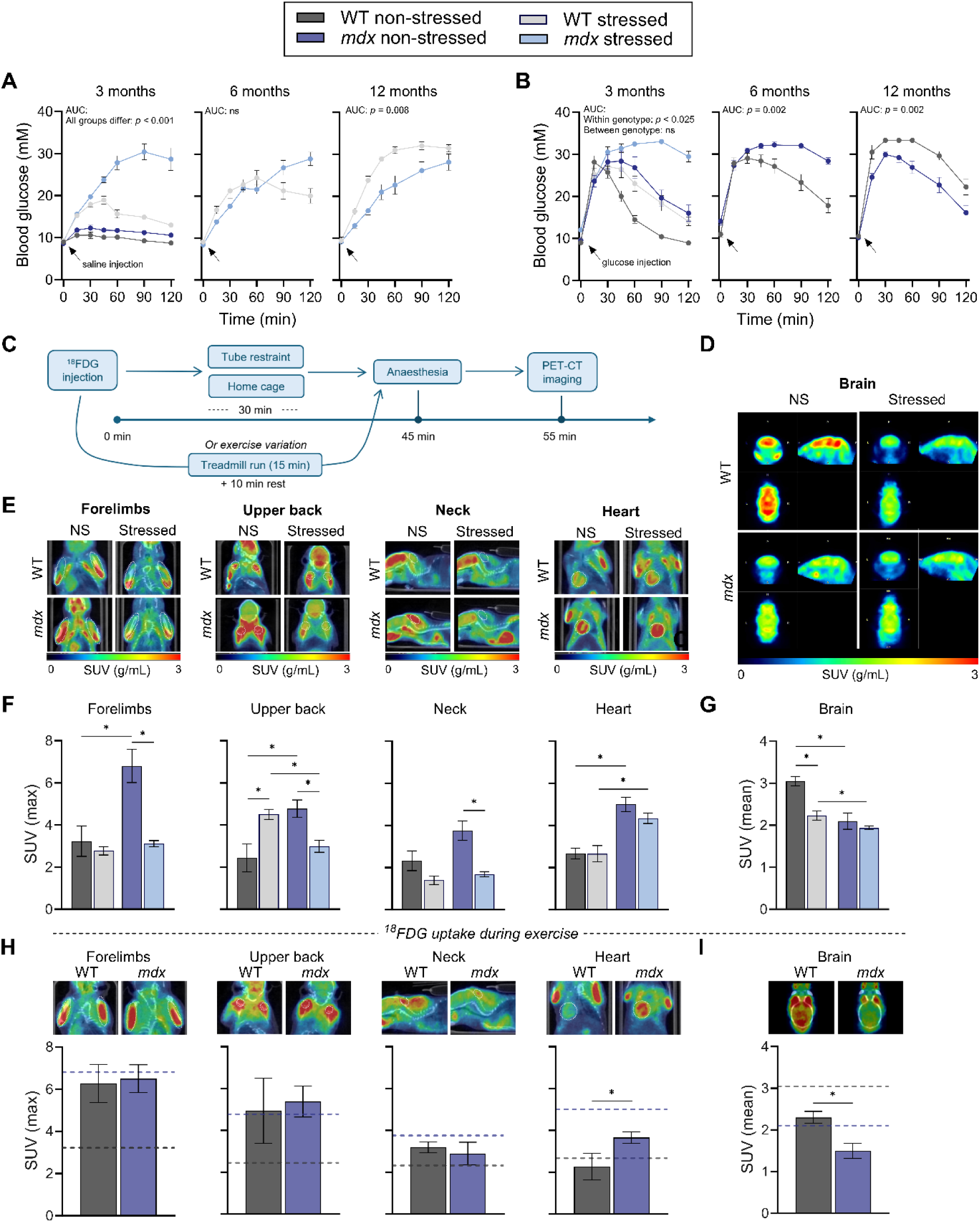
Dystrophin deficiency is associated with systemic metabolic dysfunction in *mdx* mice. Unless otherwise indicated, all measurements were performed in wild-type (WT) and *mdx* mice at rest (non-stressed) or during a tube restraint (stressed) at ∼3 months of age. **A,B,** Blood glucose concentrations at 3, 6 and 12 months of age following a 5-h fast and intraperitoneal injection of 0.9% saline (procedural control) **(A)** or 2 mg g⁻¹ glucose (glucose tolerance test (GTT)) **(B)**. Later time points (6 and 12 months) prioritised either stress responses or basal glucose handling identified at 3 months, focusing on persistence rather than re-characterisation; accordingly, non-stressed saline measurements and stressed GTTs were not performed at these timepoints. **C,** Schematic overview of the [¹⁸F] fluorodeoxyglucose (¹⁸F-FDG) uptake protocol followed by positron emission tomography-computed tomography (PET–CT) imaging to assess tissue glucose uptake under stress and exercise in mice. **D,E,** Representative PET–CT images of striated muscles (forelimbs, upper back, neck and heart) **(D)** and brain (coronal, sagittal and horizontal planes) **(E)**. **F,** Maximum ¹⁸F-FDG standardised uptake values (SUV) in striated muscles. **G,** Mean ¹⁸F-FDG SUV in the whole-brain. **H,I** Maximum ¹⁸F-FDG standardised uptake values (SUV) in striated muscles **(H)** and mean ¹⁸F-FDG SUV in the whole-brain **(I)** of mice during treadmill exercise. Dashed lines correspond to the mean baseline values, non-stressed values for WT (grey) and *mdx* (purple) from panels (F) and (G). Data were analysed using repeated-measures ANOVA, linear mixed-effects models with Bonferroni post-hoc adjustments, One-way ANOVA, Welch’s ANOVA or Kruskal-wallis. n = 4-16 per group. Data are mean ± SEM. * *P* < 0.05. Full statistical analyses and underlying raw data are provided in the Supplementary Data file.

To pinpoint tissue-level consequences of metabolic dysfunction, we used ^18^fluorodeoxyglucose (^18^FDG) uptake and positron emission tomography computer tomography (PET-CT) to map glucose uptake in the brain and striated muscle **(Fig 1C-I)**(*39*). As FDG uptake reflects integrated influences including glucoregulatory hormones, substrate availability, perfusion and stress hormones, these measurements provide an overall *in vivo* readout of tissue glucose handling. Under non-stressed conditions, *mdx* mice generally exhibited higher striated muscle ^18^FDG uptake but lower brain ^18^FDG uptake relative to WT mice **(Fig 1D-G; Fig S1F-G)**. Stress reduced brain ^18^FDG uptake in WT mice and lowered skeletal muscle ^18^FDG uptake in *mdx* mice to WT levels, while WT muscle uptake and *mdx* brain uptake remained largely unchanged **(Fig 1D-G; Fig S1F-G)**. These findings indicate that in *mdx* mice, basal glucose uptake is relatively increased in striated muscles and reduced in the brain under basal conditions, with differential responsiveness to stress across tissues.

To further probe this metabolic prioritisation, we assessed ^18^FDG uptake during treadmill exercise. Under these conditions, skeletal muscle ^18^FDG uptake did not differ between WT and *mdx* mice across forelimb, upper back and neck regions **(Fig 1F-I)**. This contrasts with the basal state, where uptake was generally elevated in *mdx* muscle, and suggests a reduced difference between genotypes during metabolic stimulation. In contrast, genotype-dependent differences persisted in the heart and brain, with *mdx* mice maintaining higher cardiac uptake and lower brain uptake relative to WT mice **(Fig 1H-I)**. These data indicate that while skeletal muscle uptake converges across genotypes during exercise, tissue-specific differences in cardiac and cerebral glucose handling are maintained.

### Altered metabolic flexibility is associated with changes in glucose handling

Because *mdx* mice exhibited impaired systemic glucose buffering, we next examined how dystrophin deficiency is associated with changes in glucose supply and metabolic flexibility across central metabolic tissues. *Mdx* mice displayed increased skeletal muscle glycogen that declined following stress; however, reduced glucose oxidation and lower glycogen phosphorylase activity at rest and following stress in *mdx* relative to WT mice are consistent with altered muscle glycogen utilisation **(Fig 2A-C)**. In contrast, liver glycogen stores were reduced at rest and after stress **(Fig 2D)**. Although hepatic glycogen phosphorylase activity increased in response to stress, hepatic glucose oxidation did not exceed WT levels, suggesting that altered systemic glucose regulation in *mdx* mice is not solely explained by increased hepatic glucose utilisation, although changes in hepatic glucose output cannot be excluded **(Fig 2D-F)**.

**Fig. 2.**
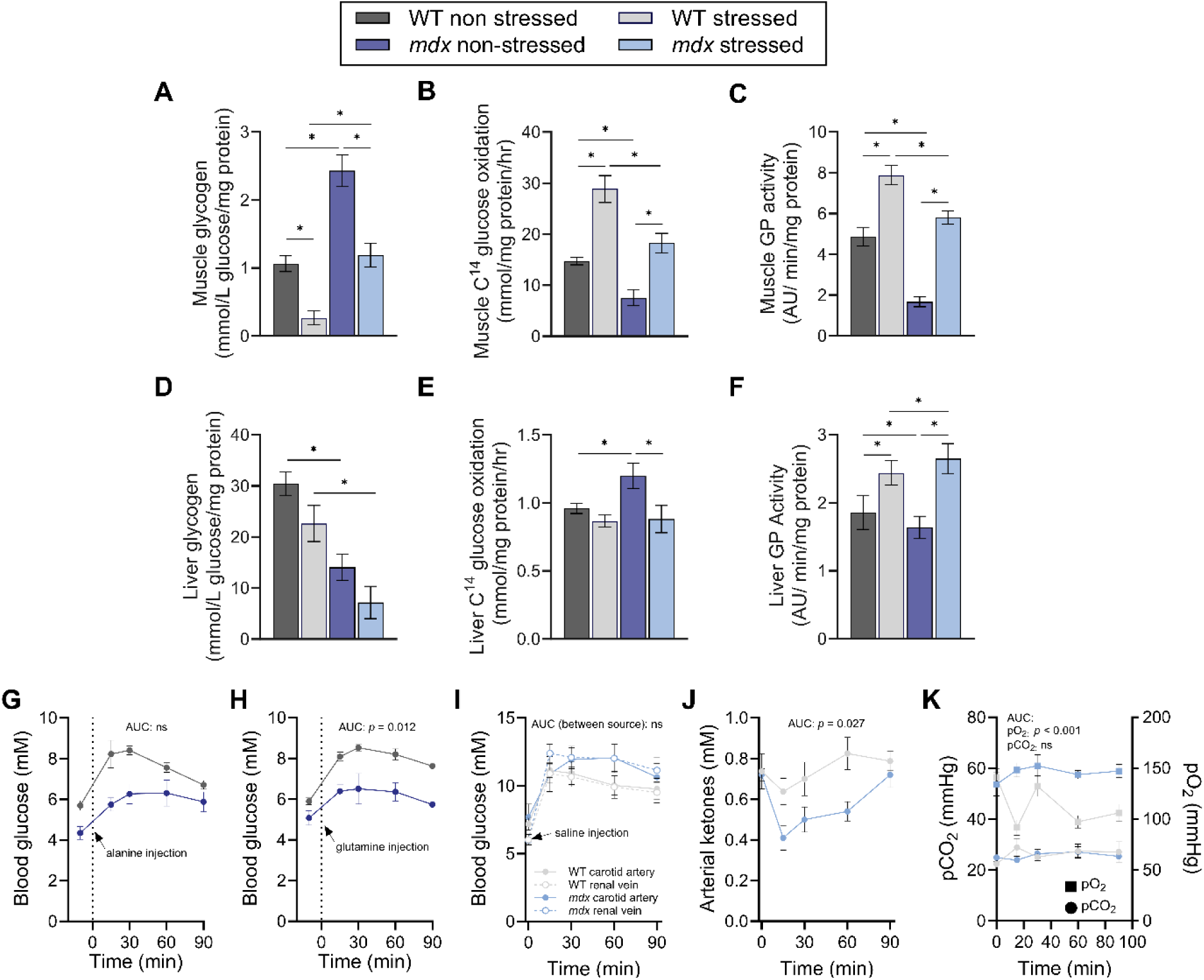
Altered substrate partitioning and gluconeogenic flux in *mdx* mice. Unless otherwise indicated, all measurements were performed in wild-type (WT) and *mdx* mice at rest (non-stressed) or during a tube restraint (stressed) at ∼3 months of age. **A,B,C,** Quadriceps muscle glycogen content **(A)**, [¹⁴C]-glucose oxidation rate **(B)**, and glycogen phosphorylase (GP) activity **(C)** measured at the end of a 2-h tube restraint (stressed) or non-stressed controls. **D,E,F,** Liver glycogen content **(D)**, [¹⁴C]-glucose oxidation rate **(E)**, and glycogen phosphorylase activity **(F)** under the same conditions. **G,H,** Blood glucose concentrations following a 5-h fast and intraperitoneal injection of alanine **(G)** or glutamine **(H)** to assess hepatic and renal gluconeogenic capacity. **I,** Blood glucose concentrations measured concurrently from the carotid artery and renal vein under stressed conditions and following a 5-h fast and 0.9% saline injection. **J,K,** Arterial ketone concentration **(J)** and partial pressure of oxygen (PO₂) and carbon dioxide (PCO₂) **(K)** measured under stressed conditions. Data were analysed using Two-way ANOVA, repeated-measures ANOVA or linear mixed-effects models with Bonferroni post-hoc adjustments. n = 6-10 per group. Data are mean ± SEM. * *P* < 0.05. Full statistical analyses and underlying raw data are provided in the Supplementary Data file.

To examine hepatic and extra-hepatic contributions to glucose homeostasis, we performed alanine (predominantly hepatic gluconeogenesis), and glutamine (extra-hepatic) tolerance tests **(Fig 2G,H)**. Glucose production from glutamine, but not alanine, was reduced in *mdx* mice, consistent with modified extra-hepatic gluconeogenic capacity **(Fig 2G,H)**. Given that the kidney can contribute to systemic extra-hepatic gluconeogenesis during stress(*40*), we assessed renal glucose output using double cannulation of the renal vein and carotid artery. Under stress, renal glucose levels did not exceed arterial levels, suggesting no net renal glucose release under these conditions **(Fig 2I)**. These findings indicate that while renal gluconeogenic capacity from glutamine is altered, renal glucose output does not appear to account for elevated systemic glucose under stress.

We next assessed alternate fuel availability and physiological stability under stress via arterial blood measurements. Arterial ketone levels were lower in *mdx* mice, consistent with differences in metabolic adaptation under stress **(Fig 2J)**. While arterial pCO₂ was comparable between genotypes, *mdx* mice exhibited elevated pO₂, suggesting differences in tissue oxygen utilisation or ventilatory-metabolic coupling **(Fig 2K)**. Despite this, arterial pH, bicarbonate, electrolytes, and oxygen-carrying capacity were preserved, indicating maintained systemic stability **(Fig S2A-G)**. Ionised calcium levels were reduced in *mdx* mice during stress, in line with altered calcium handling **(Fig S2E)**(***41***). Collectively, these data indicate reduced metabolic flexibility in *mdx* mice, characterised by altered glucose buffering and tissue-specific changes in substrate handling under stress.

### Glucose dysregulation is associated with a mismatch between signalling and insulin responses

Due to the widespread disruption of glucose buffering and metabolic inflexibility in dystrophin deficiency, we next asked whether alterations in stress-induced insulin secretion are associated with systemic metabolic vulnerability **(Fig 3A)**. To evaluate stress hormone dynamics, we measured basal and stress-induced corticosterone as an index of hypothalamic-pituitary-adrenal (HPA) axis activity. Basal levels were comparable at rest, but acute stress robustly activated the HPA axis, with an increased corticosterone response in *mdx* mice **(Fig 3B)**. Despite this heightened stress signal, *mdx* mice showed a reduced insulin response. Stress elevated circulating C-peptide, a marker of endogenous insulin secretion, in WT mice but not in *mdx* mice, resulting in lower C-peptide levels under stress **(Fig 3C)**. To distinguish potential differences in insulin action, we assessed insulin tolerance at rest and during stress. Although *mdx* mice exhibited reduced insulin sensitivity at rest, this difference was not evident under stress, where both genotypes responded similarly to insulin **(Fig 3D; Fig S3A)**. Stress altered insulin responsiveness in WT mice but not *mdx* mice, resulting in comparable responses across genotypes. These findings are consistent with a relative impairment in insulin availability under stress conditions rather than a marked change in insulin responsiveness.

**Fig. 3.**
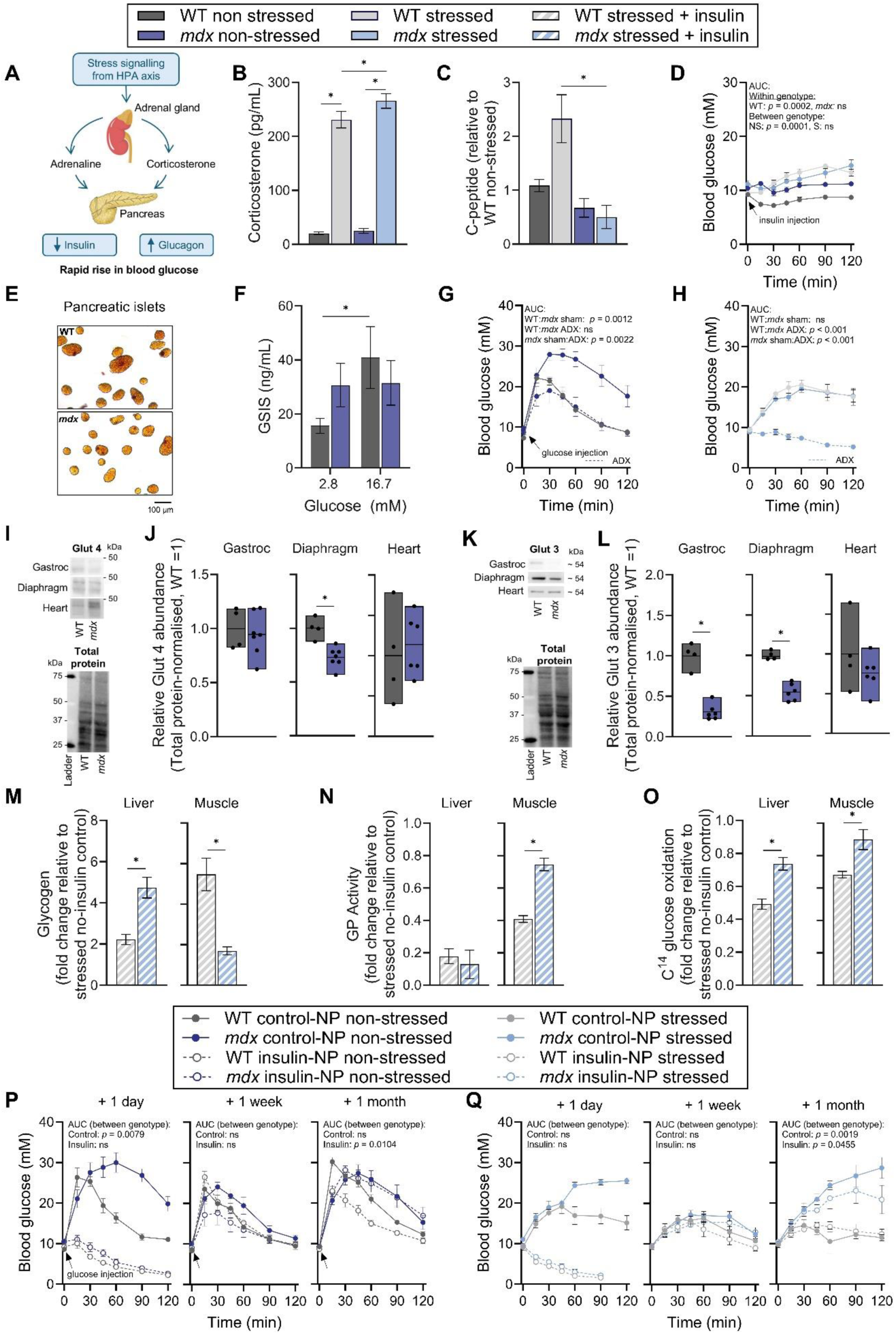
Altered coordination of stress and insulin responses in *mdx* mice. Unless otherwise indicated, all measurements were performed in wild-type (WT) and *mdx* mice at rest (non-stressed; NS) or during a tube restraint (stressed; S) at ∼3 months of age. **A,** Schematic of hypothalamic–pituitary–adrenal (HPA) axis signalling regulating pancreatic hormone secretion and systemic glucose homeostasis. **B,** Plasma corticosterone concentrations. **C,** Serum C-peptide levels, expressed relative to non-stressed WT mice. **D,** Blood glucose following a 5-h fast and intraperitoneal insulin injection (0.5 IU/kg, insulin tolerance test (ITT)). **E,** Representative pancreatic islets isolated from WT and *mdx* mice. **F,** Glucose stimulated insulin secretion (GSIS) performed from isolated islets sequentially incubated with 2.8 mM (physiological) and 16.7 mM (stimulated) glucose. **G,H,** Blood glucose concentration of adrenalectomised *mdx* mice and sham controls following a 5-h fast and intraperitoneal injection of 2 mg g⁻¹ glucose (glucose tolerance test (GTT)) **(G)**, or 0.9% saline (procedural control) **(H)**. **I-L,** Representative immunoblots **(I,K)** and quantification **(J,K)** of glucose transporter 4 (GLUT4) **(I,J)** and GLUT 3 **(K,L)** protein abundance in striated muscle extracts (gastrocnemius, diaphragm and heart), normalised to total protein and expressed relative to WT bars indicate mean with minimum–maximum values. **M-O,** Muscle and liver glycogen **(M)**, glycogen phosphorylase (GP) activity **(N)**, and [¹⁴C]-glucose oxidation rate **(O)** in stressed *mdx* mice treated with 0.5 IU/kg insulin expressed relative to non-treated controls. **L,M,** Blood glucose concentrations measured 1 day, 1 week, and 1 month after treatment with a single injection of insulin-loaded nanoparticles or control nanoparticles following a 5-h fast and intraperitoneal injection of 2 mg g⁻¹ glucose (GTT) **(P)**, or 0.9% saline (procedural control) **(Q)**. Data were analysed using One-way ANOVA, Two-way ANOVA, repeated-measures ANOVA or linear mixed-effects models with Bonferroni post-hoc adjustments. n = 4-10 per group. Data are mean ± SEM (unless otherwise stated). * *P* < 0.05. Full statistical analyses and underlying raw data are provided in the Supplementary Data file.

The pancreas-stomach interface has been implicated in dystrophin-deficient animals (*mdx*4cv) with downregulation of dystrophin and DGC components(*42*). To examine pancreatic function, we measured glucose-stimulated insulin secretion in isolated pancreatic islets. Islets from *mdx* mice displayed normal basal insulin release but a reduced response to high glucose concentrations **(Fig 3E-F, Fig S3B-C)**. This occurred despite equivalent islet size **(Fig 3E-F, Fig S3B-C)**. These observations indicate that dystrophin deficiency is associated with functional changes in metabolic tissues other than muscle, including the pancreatic islets. Finally, to assess the contribution of stress hormone signalling to glucose dynamics, we performed adrenalectomy (ADX) and subsequent glucose handling measurements. Removal of adrenal stress hormones improved glucose tolerance and attenuated stress-induced hyperglycaemia in *mdx* mice, demonstrating that adrenal hormones contribute to the observed metabolic phenotype **(Fig 3G-H)**. Together, these data are consistent with altered coordination between stress signalling and insulin secretion in *mdx* mice, which may contribute to impaired metabolic regulation under stress.

Having established altered coordination between stress signalling and insulin secretion, we next examined how these endocrine changes are associated with tissue-level glucose uptake and storage, and whether increasing insulin availability could modify these responses. Glucose entry into muscle and heart is largely mediated by the insulin-responsive transporter GLUT4(*43*), whereas GLUT3 supports basal, insulin-independent uptake, particularly in neurons(*44*). GLUT4 protein abundance was similar between genotypes in gastrocnemius and heart but reduced in the diaphragm of *mdx* mice **(Fig 3I-J; Fig S4)**. However, prior studies suggest that diaphragm-specific reductions in *mdx* mice may in part reflect increased fibrosis, which can dilute muscle-specific proteins in tissue lysates(*45*). Across skeletal muscles, GLUT3 protein abundance was decreased in *mdx* mice, whereas levels in the heart and brain were comparable between genotypes **(Fig 3K–L; Fig S3D-F; Fig S5)**. Reduced skeletal muscle GLUT3 may influence basal or stress-associated glucose uptake, although this occurs alongside elevated ^18^FDG uptake in unstressed *mdx* muscle, indicating that multiple factors contribute to *in vivo* glucose handling. Together, these data indicate that while total GLUT4 abundance is largely preserved in key muscles, differences in GLUT3, a key neuronal receptor, may also contribute altered glucose regulation in dystrophin deficiency.

We next assessed how these differences are reflected in glycogen storage and glucose oxidation, and whether transient insulin supplementation modifies these responses. Insulin administered prior to stress, expressed as relative change from no-insulin controls, increased hepatic and skeletal muscle glycogen accumulation in both genotypes (>1), with a greater increase observed in *mdx* liver and comparatively lower increase in *mdx* muscle, relative to WT tissues **(Fig 3M)**. While overall depleted relative to the no-insulin controls (<1), muscle glycogen phosphorylase activity remained higher in *mdx* mice relative to WT mice, with no genotypic differences in hepatic phosphorylase activity **(Fig 3N)**. Insulin-mediated suppression of glucose oxidation was reduced in *mdx* skeletal muscle and liver compared to WT tissues **(Fig 3O)**. These observations indicate that glycogen storage and glucose utilisation respond differently across tissues in *mdx* mice under insulin-stimulated conditions, supporting previous observations(*29*).

To assess whether increasing insulin availability could modify systemic glucose control, we administered glucose-responsive insulin nanoparticles(*46*). These allow insulin release in response to circulating glucose and provide controlled delivery under dynamic conditions. Treatment reduced blood glucose excursions during glucose tolerance testing and attenuated stress-induced hyperglycaemia in *mdx* mice **(Fig 3P,Q).** Effects were most evident one day post-administration, diminished by one week (likely reflecting residual nanoparticle activity), and were no longer detectable at one month **(Fig 3P,Q)**. No overt tissue toxicity was observed **(Fig S6A-B).** Collectively, these findings indicate that tissue glucose handling in dystrophin deficiency reflects the combined influence of altered endocrine responses and tissue-specific metabolic context. Increasing insulin availability can transiently improve systemic glucose control, highlighting a modifiable component of the metabolic phenotype.

### Insulin improves systemic metabolic responses under stress

To further examine how insulin availability associates with tissue-specific glucose handling and systemic metabolic responses to stress, we quantified ^18^FDG uptake in striated muscles and brain and monitored whole-body energy expenditure in insulin-treated animals under basal and acute stress conditions **(Fig 4A-B)**. At rest, insulin affected ^18^FDG uptake in skeletal muscle and brain in both genotypes equally, consistent with retained responsiveness of *mdx* mice to insulin *in vivo*, whereas cardiac uptake remained elevated in *mdx* mice relative to WT mice **(Fig 4A-D, Fig S3G)**. Under stress, skeletal muscle uptake remained largely stable, although *mdx* mice showed lower uptake than WT, while cardiac ^18^FDG remained elevated **(Fig 4A,C)**. Brain uptake declined under stress in both genotypes, indicating a shared stress response **(Fig 4B,D, Fig S3G)**. Whole-body metabolic parameters were assessed following an acute scruff-restraint stress exposure, aligning with previous stress literature in *mdx* mice(*33*, *34*, *47*). Insulin treatment was associated with increased post-stress physical activity and normalised *V̇*O₂, RER and heat production in *mdx* mice **(Fig 4E-L)**. These measurements capture responses to a short stress challenge, but are consistent with improved metabolic responsiveness in insulin-treated animals. Collectively, these data indicate that increasing insulin availability modulates tissue glucose uptake and systemic metabolic and behavioural responses in *mdx* mice under stress.

**Fig. 4.**
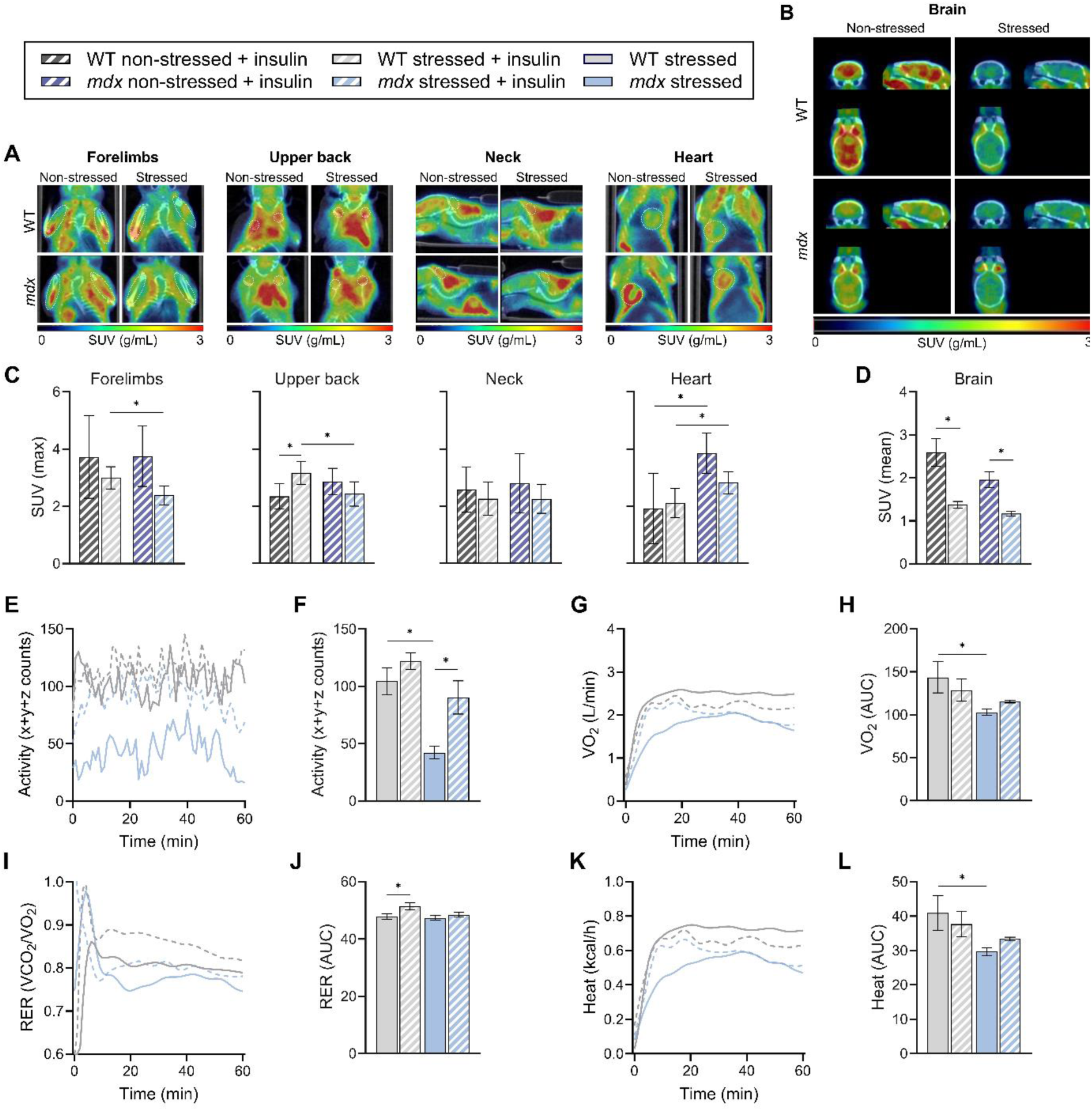
Insulin modulates stress-induced glucose uptake and whole-body metabolism. Unless otherwise indicated, all measurements were performed in wild-type (WT) and *mdx* mice at rest (non-stressed) or during a tube restraint (stressed) at ∼3 months of age. **A,B,** Representative positron emission tomography–computed tomography (PET–CT) images of striated muscles (forelimbs, upper back, neck and heart) **(A)** and brain (coronal, sagittal and horizontal planes) **(B)** from mice treated with exogeneous insulin prior to stress or non-stress conditions. **C,** Maximum [¹⁸F]fluorodeoxyglucose (¹⁸F-FDG) standardised uptake values (SUV) in striated muscles. **D,** Mean ¹⁸F-FDG SUV in the whole-brain. **E-L,** Physical activity (counts per minute) **(E)**, mean physical activity (counts per minute) **(F)** oxygen consumption (VO_2_) over time **(G)**, oxygen consumption (VO_2_) area under the curve (AUC) **(H),** respiratory exchange ratio (RER) over time **(I)**, RER AUC **(J)**, heat production **(K)**, and heat AUC **(L))** measured in Promethion metabolic cages following intraperitoneal injection with 0.9% saline or 0.5 IU/kg insulin prior to a 30 s scruff-restraint. Data were analysed using repeated-measures ANOVA or linear mixed-effects models with Bonferroni post-hoc adjustments. n = 5-8 per group. Data are mean ± SEM. * *P* < 0.05. Full statistical analyses and underlying raw data are provided in the Supplementary Data file.

### Glucose offloading improves systemic glucose control and muscle function

Having identified altered coordination of stress and insulin responses in dystrophin-deficiency, we next examined whether reducing circulating glucose via the SGLT2 inhibitor empagliflozin (EMPA) could improve systemic metabolic control. EMPA was compared with prednisolone, a current standard of care, and their combination to assess effects on glucose regulation and muscle function. EMPA reduced stress-induced glucose excursions in *mdx* mice, bringing responses closer to WT levels **(Fig 5A).** This effect was selective, as EMPA had minimal impacts on fasting glycaemia, in contrast to prednisolone, which lowered baseline glucose levels **(Fig 5B)**. These effects were accompanied by increased urinary glucose excretion in EMPA-treated animals, consistent with enhanced renal glucose clearance **(Fig 5C)**. Circulating ketone levels were not increased **(Fig 5D)**, indicating no overt shift toward ketone-dominated metabolism under these conditions.

**Fig. 5.**
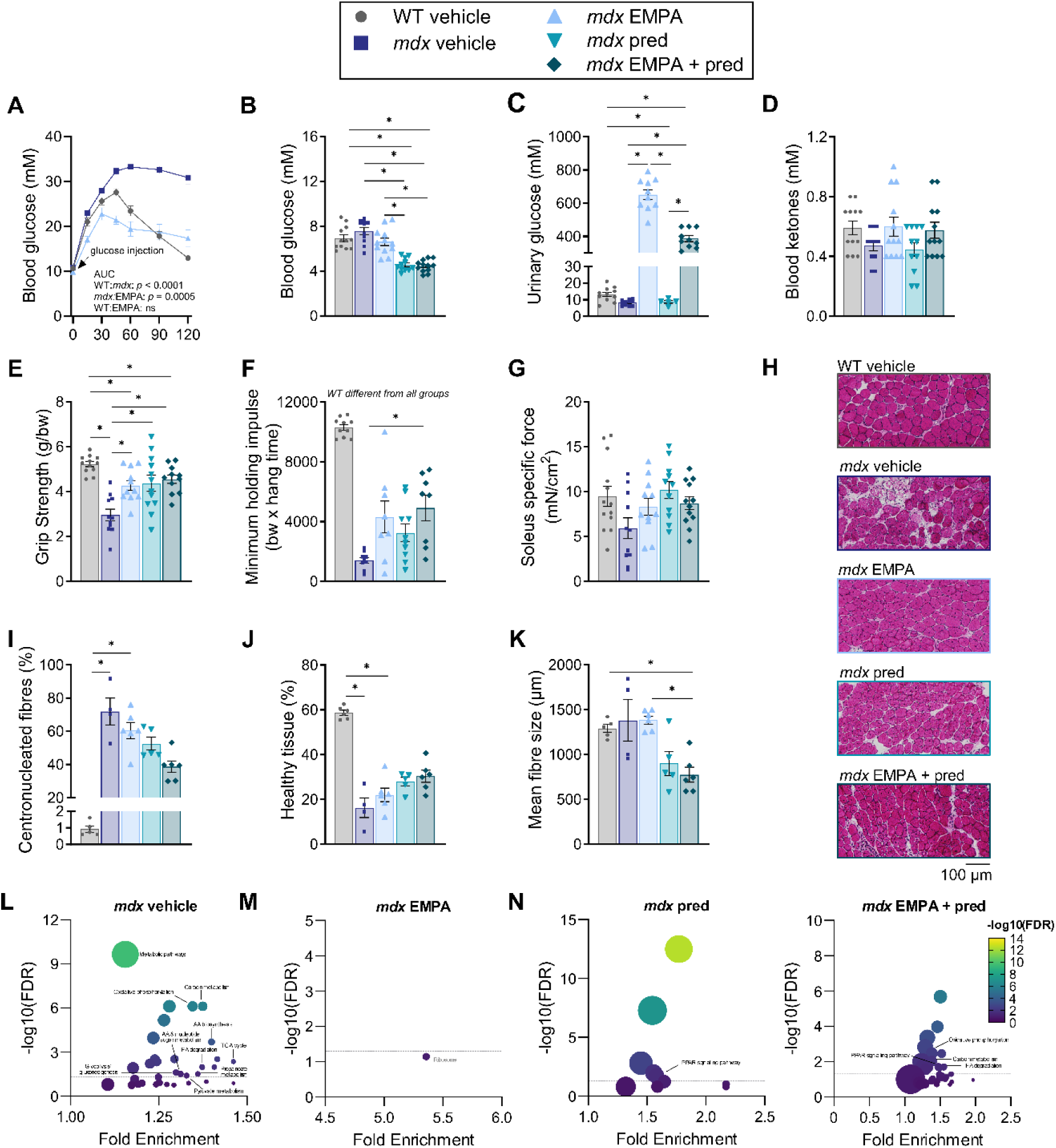
Glucose offloading with empagliflozin (EMPA) improves stress-associated glucose responses and muscle function in *mdx* mice. A, Blood glucose concentration under acute stress following a 5-h fast and intraperitoneal injection of 2 mg g⁻¹ glucose (glucose tolerance test (GTT))**. B**, Endpoint fasting blood glucose levels. **C**, Urinary glucose excretion. **D**, Fasted circulating blood ketone concentrations. **E**, Forelimb grip strength. **F**, Four-limb wire hang performance expressed as minimum holding impulse. **G**, Soleus specific force. **H**, Representative haematoxylin and eosin–stained cross-sections of soleus muscle. **I**, Percentage of centrally nucleated fibres. **J**, Percentage of healthy tissue area. **K**, Mean myofibre cross-sectional area. **L-O**, KEGG pathway enrichment analysis of muscle proteomics comparing **(L)** *mdx* vehicle relative to WT vehicle control, and **(M)** EMPA, **(N)** prednisolone, and **(O)** EMPA + prednisolone relative to *mdx* vehicle control. Fold enrichment is plotted against −log₁₀(FDR), with dot size proportional to the number of hits per pathway; the horizontal dotted line denotes the significance threshold (FDR = 0.05). Only metabolic pathways are labelled for clarity except, in (m) where the only enriched non-metabolic pathway is labelled in grey. Data were analysed using Kruskal-Wallis with Dunn’s post-hoc or Welch’s ANOVA and proteomics data were analysed using TMT analyst from Monash Proteomics and Metabolomics Platform with subsequently identified significant proteins uploaded into ShinyGO. *n* = 4-12 per group. Data are mean ± SEM. * *P* < 0.05. Full statistical analyses and underlying raw data are provided in the Supplementary Data file.

To assess whether systemic metabolic changes and muscle function could be improved in parallel, we evaluated muscle performance. EMPA treatment improved forelimb grip strength and holding impulse in *mdx* mice to a similar extent as prednisolone, although neither intervention improved isolated soleus-specific force **(Fig 5E-G)**. These functional improvements occurred without detectable changes in overall muscle architecture, including centronucleation, healthy tissue area, or myofibre size **(Fig 5H–K)**. Consistent with this, EMPA did not exacerbate features consistent with myofibre atrophy, in contrast to prednisolone-treated muscles, which exhibited reduced fibre size **(Fig 5H,K)**.

To evaluate how glucose offloading influences tissue responses under these conditions, we examined skeletal muscle molecular profiles. Given that previous studies have primarily focused on immunomodulatory, extracellular matrix, and cytokine signalling pathways in dystrophin deficiency, we specifically interrogated metabolism-related processes to complement these findings(*48*). Pathway enrichment analysis mapped proteomic data to KEGG pathways using ShinyGO and direct focus was given to significant metabolism-related terms **(Fig 5L-O)**. WT and *mdx* gastrocnemius muscle showed differences in metabolic pathways, including glycolysis/gluconeogenesis and mitochondrial respiration **(Fig 5L-O, Fig S7A,H-L, Table S1-S4)**. EMPA treatment was not associated with large-scale shifts in these pathway enrichments, whereas prednisolone and combination treatment were associated with changes in pathways linked to oxidative phosphorylation, carbon metabolism and fatty acid degradation **(Fig 5N–O**; see **Fig. S7** for pathway directionality). As enrichment analysis reflects changes in protein abundance rather than pathway activity, these findings suggest that EMPA primarily influences systemic glucose handling, with limited detectable alteration of muscle metabolic protein profiles under these conditions. Together, these data indicate that reducing circulating glucose via renal excretion improves stress-associated glucose regulation and is accompanied by improved muscle function in *mdx* mice. These effects occur in the absence of major changes in muscle structure or broad proteomic pathway shifts, suggesting that systemic metabolic context contributes to functional outcomes in dystrophin deficiency.

## DISCUSSION

Dystrophin deficiency reveals an underappreciated vulnerability in systemic metabolic control beyond its structural role in muscle. Heightened stress-induced hyperglycaemia and glucose intolerance in *mdx* mice are not readily explained by intrinsic peripheral tissue defects alone, but are associated with altered coordination between insulin secretion and stress responses. This distinction is important given ongoing controversy regarding glucose intolerance in *mdx* mice, with prior studies variably reporting preserved glucose clearance(*49*), delayed glycaemic peaks without differences in area under the curve(*29*), and elevated respiratory exchange ratios indicative of altered substrate use(*29*, *50*). In the present study, physiological stress is accompanied by a marked increase in glucocorticoid signalling, whereas insulin secretion does not increase to the same extent, consistent with a relative mismatch in endocrine responses. This imbalance emerges in a hypermetabolic system, as *mdx* mice exhibit elevated oxygen consumption, increased whole-body energy expenditure, and a tendency toward carbohydrate utilisation(*50*), indicative of heightened metabolic demand.

These findings point to constraints in metabolic flexibility rather than absolute substrate availability. Dystrophin-deficient skeletal muscle exhibits elevated basal glucose uptake and increased glycogen storage, alongside reduced glucose oxidation and a limited increase in uptake during exercise, in line with reduced metabolic adaptability under challenge. Liver glycogen stores are reduced but do not appear to fully account for stress-induced hyperglycaemia, aligning with previous reports of reduced hepatic glycogen content without major alterations in overall glycogen metabolism(*29*). This pattern could reflect the elevated energetic demands described in dystrophin-deficient muscle, which sustains greater metabolic demand despite chronically reduced resting ATP levels imposed by calcium buffering (e.g., stress-induced ionised calcium drop shown here), membrane repair, and regenerative turnover(*21*, *50–52*). At the level of glucose transport, total GLUT4 abundance is largely preserved, whereas GLUT3, a key neuronal glucose receptor, is reduced in skeletal muscle but maintained in the brain, suggesting potential differences in basal or insulin-independent glucose uptake across tissues. However, *mdx* mice exhibited ∼30% lower baseline brain ¹⁸FDG uptake relative to WT mice and showed limited further decline under stress, suggesting that factors such as substrate availability or perfusion may constrain cerebral glucose handling despite preserved GLUT expression. This aligns with evidence of reduced free glucose but increased oxidative ¹³C-glucose flux in the *mdx* brain(*12*), and reinforces a dystrophin-related neuronal metabolic defect rather than exclusive peripheral diversion.

Increasing insulin availability partially mitigates these limitations. *Mdx* mice maintain basal insulin secretion but show a blunted increase in C-peptide during stress, and isolated islets demonstrated attenuated glucose-stimulated insulin secretion. While insulin tolerance tests suggest reduced insulin sensitivity at baseline, this difference is diminished under stress, where both genotypes show a comparable response to insulin, consistent with a predominant role for insulin availability under stress conditions. Exogenous insulin increases glucose uptake and improves systemic metabolic responses, including normalisation of stress-induced physical inactivity - a hallmark behavioural phenotype(*30*). These observations support a model in which altered insulin dynamics contribute to stress-associated metabolic dysregulation, rather than reflecting primary intrinsic insulin resistance. Indeed, insulin resistance reported in DMD during disease progression may more likely arise from secondary factors such as reduced ambulation, obesity, or chronic glucocorticoid exposure(*8*). In addition, given that glucoregulatory hormones are strongly stress-responsive and neurally regulated(*53*), and that dystrophin is also expressed in the brain, the altered brain ^18^FDG uptake and HPA axis activation observed here are consistent with a contribution from central dystrophin involvement to the systemic metabolic phenotype.

Targeting circulating glucose levels provides an additional means of modifying stress-induced hyperglycaemia. Glucose-responsive insulin-loaded nanoparticles improved glucose control transiently, indicating that glucose-responsive insulin delivery can partially recouple insulin availability with dynamic metabolic demand under stress conditions. Similarly, EMPA reduced stress-induced glucose spikes without substantially altering fasting glycaemia, consistent with increased renal glucose excretion and reduced reliance on tissue glucose disposal. Circulating ketone levels were not increased, indicating that these effects occurred without a detectable shift toward ketone-dominated metabolism. Notably, EMPA treatment was associated with improved muscle function, as seen by a ∼45% increase in forelimb grip strength, despite minimal changes in muscle structure of broad proteomic pathways. These findings suggest that modulation of systemic glucose handling and improvements in muscle function can occur in parallel, potentially reflecting complementary rather than directly coupled effects. Supporting recent studies reporting functional improvements via anti-fibrotic or muscle-intrinsic remodelling pathways(*48*), these data indicate that systemic metabolic interventions may offer an additional route to enhance functional outcomes. This raises the possibility that targeting systemic glucose regulation alongside muscle pathology may provide a dual approach to treatment in dystrophin deficiency.

Together, these findings help reconcile conflicting reports of glucose intolerance in DMD(*29*, *49*, *50*, *54*) and support the view that metabolic abnormalities are most evident under physiological challenge. Rather than reflecting a primary defect in glucose uptake or insulin action, the phenotype is consistent with altered coordination between endocrine responses and tissue-level metabolic demands. Tissue-specific patterns, including relatively preserved cardiac uptake and reduced cerebral uptake, further point to differential constraints across organ systems in dystrophin deficiency. Interventions that either increase insulin availability or reduce circulating glucose levels can improve systemic metabolic responses and muscle function without structural remodelling. These observations highlight the importance of systemic metabolic regulation in shaping functional outcomes and suggest that targeting whole-body glucose handling may represent a complementary therapeutic strategy in dystrophin deficiency. Further investigation of how central (neural) and peripheral (muscle) control mechanisms interact to coordinate these responses will be important to define how neural and muscle-specific processes jointly shape systemic metabolic responses under stress. Together, these findings highlight dystrophin deficiency as a multi-system disorder in which disrupted coordination between neural, endocrine and muscle metabolism limits adaptive responses to stress.

### Study Limitations

These findings should be interpreted in the context of the inherent complexity of metabolic regulation under stress. Coordination between endocrine signalling, including insulin and other glucoregulatory hormones, tissue metabolism, and central control operates across multiple scales and cannot be fully resolved within a single experimental paradigm or by discrete temporal sampling. However, the approach used here captures key features of this coordinated response and supports a broader interpretation of the phenotype. This provides a basis for future studies to interrogate tissue-specific metabolic pathways, distinguish systemic from cell-intrinsic contributions, and define the role of neural circuits in shaping metabolic responses. While the present data demonstrate that this phenotype can be modulated experimentally, further work across longer timeframes and in models that more closely reflect human disease will be required to determine how these mechanisms translate to clinical settings.

## MATERIALS AND METHODS

### Ethics statement

All experiments were performed with ethical approval from the Monash University (38106, 21897, 41056, 34696, E/8220/2027/M), Virginia Tech University, the Vanderbilt University (DK135073, DK020593) and Victoria University (AEETH 22-005) Animal Ethics Committees. All animals were housed and treated in accordance with the Animal Welfare Act and the ARRIVE guidelines for animal experimentation(*55*).

### Experimental mice

Male C57BL/10ScSn/Ozarc or C57BL/10ScSn/J (wildtype; WT) and C57BL/10ScSn-*Dmd^mdx^*/Ozarc or C57BL/10ScSn-*Dmd^mdx^*/J (*mdx*) mice were purchased from Animal Resources Centre (Ozgene ARC; Perth, WA, Australia) or Jackson Laboratories (Bar Harbor, Maine, United States) respectively, and allowed to acclimate for one week before commencing any experiments. The holding rooms were maintained on a 12/12 h light/dark cycle with humidity between 40 - 70% and temperature at 21°C. Food and water were provided *ad libitum* and the welfare of the animals were monitored daily.

### Metabolic phenotyping studies

For standard blood glucose assessments and glucose tolerance tests, mice were fasted for ∼5 h and for the insulin tolerance test mice were fasted for ∼4 h (beginning at 0700-0900 h; water provided *ad libitum*). After 4 - 5 h of fasting, body weight was measured. A clean scalpel blade was used to make a cut (snip; < 2 mm) at the very end of the tail where all veins meet. Mice were placed in tube restrainers for 2 h to obtain blood samples from animals in a ‘stressed state’. Prolonged tube restraint (2 h) was used in metabolic phenotyping experiments to enable continuous assessment of glucose and hormonal responses under sustained stress. For animals in an unstressed state, they were quickly handled by the tail, blood sampled and released back into their home cage. Water was then removed. Blood was collected by gentle manipulation/massaging of the tail from the base to tail tip where the snip was performed. Baseline blood glucose levels were measured using a standard handheld glucometer (2-5 µL of blood is required per measurement). After the baseline blood glucose measurement was obtained, D-glucose (2 mg of 50% glucose/g body mass) for the glucose tolerance test (or saline for the sham glucose tolerance test) or insulin (insulin 100 IU diluted to 0.20 IU making a 500-fold solution, 0.50 IU insulin/g body mass) for the insulin tolerance test was administered via intraperitoneal injection. Blood samples were collected to measure blood glucose levels at 15, 30, 45, 60, 90, and 120 minutes following the intraperitoneal injection. After the 120-minute blood sample, mice were returned to their home cage and given free access to food and water.

### 18FDG uptake studies

To map brain and striated tissue activity under stress ^18^FDG uptake was assessed using PET/CT scanning following no-stress or tube-restraint stressor. All mice were injected with ∼15 MBq of ^18^FDG in the intraperitoneal space with a 30 G needle. Non-stressed (control): Mice were anaesthetised with isoflurane (1 – 2%) in 100% oxygen, and when no longer mobile injected with ^18^FDG and placed back in their home cage for 30 min prior to imaging. Stressed (Tube-restraint): Mice were injected with ^18^FDG placed into a plexiglass tube-restraint for 30 min prior to imaging. For exercise-induced uptake experiments mice were acclimatised on a treadmill for 5 mins, three times a week for 3-weeks prior to the study. Mice were fasted for 18h then injected with ∼15 MBq of ^18^FDG as described above before being run on a treadmill at 10m/min for 15 mins, followed by 10 mins of rest in the home cage and then imaging. For insulin treatment experiments, mice were injected with 0.5 IU/g of human insulin in the intraperitoneal space 1 minute prior to the ^18^FDG injection. Basal, exercise and insulin ^18^FDG studies were conducted in separate cohorts; therefore, comparisons reflect between-group analyses rather than longitudinal within-animal changes.

### PET-CT scanning and data analysis

At 45 min post injection all mice were anesthetized with 4% isoflurane in 100% oxygen and readied for PET scanning, which commenced at 55 min post injection for 10 min using a Mediso Nanoscan PET-CT small animal scanner (Mediso, Budapest, Hungary). Animals were scanned 2-4 at a time and PET data was collected using a static acquisition protocol with decay correction set to the acquisition midpoint. PET data was reconstructed using the Teratomo3D mode with 4 iterations and 6 subsets at an isometric voxel size of 0.4mm. CT data was scanned using a ZigZag protocol with 360 projections at 50 kVp and 0.67 mAmp with an exposure time of 170 ms. Reconstruction used a Feldkamp filtered back projection algorithm and generated an isometric voxel size of 0.125 mm.

PET images were analysed using PMOD (PMOD LLC, Zurich, Switzerland). Mice were individually cropped, and radioactive dose data manually entered. An identical head region was cropped for the PET and CT images, the head CT image was registered to the Ma_Benveniste_Mirrione CT brain template using deform matching in the PFUS software (**Fig. S1f**). The result was manually checked, and the resulting transformation was applied to the PET head image. Brain volumes of interest (VOIs) from the Ma_Benveniste_Mirrione template were used to quantitate uptake in specific brain regions. For muscle regions, manually drawn VOIs were placed over the scapula, forelimb and neck muscles. Cardiac ventricular uptake was determined by placing a sphere over the cardiac region and thresholding to 55% of the maximal SUV value to isolate the left ventricle uptake.

### [^14^C] Glucose oxidation assay

The liver and the quadriceps were surgically dissected. Approximately 15-50 mg of tissue was placed in 200 μl of modified sucrose EDTA medium (SET buffer) containing 250 mM sucrose, 1 mM EDTA, 10 mM Tris-HCl, and 1 mM ATP at pH 7.4. Tissues were minced with scissors, then SET buffer was added to create an approximate 20-fold dilution (weight: volume). Samples were homogenised in a Potter-Elvehjem glass homogenizer for 10 passes over 30s at 150 rpm with a Teflon pestle and then incubated in U— 14 C-glucose (American Radiolabeled Chemicals, St. Louis, MO, USA). Production of 14-CO_2_ was assessed by scintillation counting (Beckman Coulter 4500) and used to estimate glucose oxidative capacity. Total protein content in tissue homogenates was measured via BCA (Thermo Fisher Scientific, Waltham, MA, USA) and used to normalise oxidation values.

### Glycogen Phosphorylase Assay

Glycogen phosphorylase activity was performed using the Abcam Glycogen Phosphorylase Assay Kit (Abcam) according to the manufacturer’s instructions. Samples were measured in duplicate.

### Glycogen levels

Liver and muscle samples were collected from control and *mdx* mice. Liver and quadricep samples were lysed in 10 mM sodium acetate buffer (pH 4.6). Amylo-1-6-glucosidase was added to tissue lysate and incubated at 37°C for 2 h to hydrolyse glucose from glycogen. Samples were centrifuged (10,000 RPM) at room temperature, and the supernatant was analysed using the Glucose (GO) assay kit (Sigma) following the manufacturer’s protocol. Samples without amylo-1-6-glucosidase were processed similarly to the cell samples and subtracted from the results as a control for free cellular glucose. Tissue lysates were analysed for protein content (BCA assay), and results were normalised to total protein. Samples were processed identically across groups, and glycogen-derived glucose measurements were used for relative comparisons between conditions.

### Alanine and glutamine tolerance tests

Tolerance tests were performed by the Vanderbilt Mouse Metabolic Phenotyping Center. Mice were fasted for 18 h. A baseline blood glucose measurement was performed (t=0 min) from tail blood, immediately followed by administration of a L-glutamine or L-alanine solution (2 g/kg) by intraperitoneal injection. Tail blood glucose was measured at 15, 30, 60, and 90 min.

### Double cannulation and stress experiments

Double cannulation surgeries and subsequent restraint tests were performed by the Vanderbilt Mouse Metabolic Phenotyping Center. One day before restraint testing, mice were anesthetised with isoflurane and given ketoprofen (5 mg/kg, s.c.) for analgesia. The carotid artery was catheterised as described previously(*56*). For renal catheteridation, a midline abdominal incision was made, and the cecum was gently exteriorized and kept moist with sterile saline-soaked gauze. After exposing the gonadal and renal veins, two silk sutures were placed beneath the gonadal vein; the caudal suture was ligated, and a sterile catheter (Instech) was inserted and advanced 4–6 mm to the junction of the renal vein. The catheter was secured with both sutures, the abdominal wall was closed, and the catheter was tunneled subcutaneously to the interscapular site. Carotid and renal catheters were connected to a two-channel Vascular Access Button (Instech), and both lines were flushed and locked with heparinised saline (200 U/mL).

The day after surgery, mice were fasted for 4 h before undergoing a restraint stress test. The vascular access button was flushed, connected to external lines for remote renal and arterial blood sampling, and baseline glucose was measured from both catheters, followed immediately by intraperitoneal saline injection (10 mL kg⁻¹) and restrainer placement (t = 0 min). Renal glucose (Contour Next) and arterial ketones (Precision Xtra) were measured at 15, 30, 60 and 90 min. In one cohort, arterial iSTAT (CG8+, Abbott) measurements were performed at 0 and 30 min, with arterial glucose measured at 15, 60 and 90 min. In a second cohort, arterial iSTAT measurements were performed at 15, 60 and 90 min, with arterial glucose measured at 0 and 30 min. Mice were euthanised after the final sample.

### Stress adaptation studies

Mice were placed in a tube restrainer for five minutes with blood glucose levels measured at the start and end of each tube-restraint stressor as described above. This was repeated every 30 minutes over 5 h (10 stress insults in total). Standard GTTs were performed as described above, one day prior, and one day, one week and one month following the repeated stress insults.

### C-Peptide and corticosterone ELISA

Serum levels of C-peptide were measured using C-Peptide ELISA kit from ALPCO (ALPCO, NH). In brief, serum was collected, and samples were prepared according to the manufacturer’s protocol. C-peptide concentrations were measured in triplicate, with a common standard used across assay plates. Corticosterone ELISA was measured using Mouse/Rat Corticosterone ELISA kit from ALPCO (ALPCO, NH). Quantitative determination of corticosterone in mouse serum was performed according to the manufacturer’s protocol. Samples were measured in duplicate.

### Islet isolation and glucose-stimulated-insulin-secretion studies

Pancreatic islets were isolated from mice at 10 weeks of age by collagenase digestion at 37°C using 1.4 mg/mL Collagenase-P in Hanks Buffered Salt Solution supplemented with 4.2 mM NaHCO_3_, 0.1% BSA and 20 ug/mL DNAse. The islets were purified using a 1.100 g/mL Histopaque gradient, handpicked under an inverted microscope and cultured in RPMI 1640 + L-glutamine supplemented with 10% FBS and 1x penicillin/streptomycin overnight at 37°C. Islets were cultured for 24-48 h and bright field images were taken for quantification of islet size using ImageJ.

Islet insulin secretion was measured in modified Krebs-Ringer Bicarbonate-Hepes Buffer Solution (KRBH) on ten handpicked islets per mouse. Islets were incubated with KRBH containing 2.8 M glucose (basal glucose) for 1 h at 37°C to stabilize cultures and wash away media. Then the islets were incubated in basal glucose (2.8 mM) in KRBH for 1 h, followed by stimulatory glucose (16.7 mM) in KRBH. Supernatants were collected for both basal and stimulatory glucose and stored at −20°C. The insulin concentration in the supernatant was assayed using the ultrasensitive mouse insulin ELISA kit (ALPCO), in accordance with the manufacturer’s guidelines.

### Adrenalectomy surgery

Bilateral adrenalectomy was performed under general anaesthesia using a lateral approach. Small flank incisions were made between the last rib and iliac crest, the abdominal wall was opened, and the adrenal glands were identified at the cranial pole of the kidneys and excised. Incisions were closed using standard absorbable sutures and post-operative analgesia was provided for 48 h post-surgery. Sham-operated controls underwent identical surgical exposure without gland removal.

### Western blotting of glucose transporters

Brain and muscle tissues (heart, gastrocnemius, diaphragm, and triceps) were snap-frozen in chilled isopentane and homogenized in liquid nitrogen. Lysates were prepared in RIPA buffer containing protease and phosphatase inhibitors (Pierce) during a 1 h rotation at 4°C. Following protein quantification (BCA assay), 20 μg of lysate were separated on Criterion TGX gels and transferred to PVDF membranes (100 V, 1.5 h).

After blocking in 5% BSA or skim milk in TBST for 1 h, membranes were incubated overnight at 4°C with primary antibodies for GLUT3 (ProteinTech, 1:500) or GLUT4 (Thermo Fisher, 1:500). Detection was performed with a goat anti-rabbit Alexa Fluor 594 nm secondary antibody (1:10,000, 1.5 h). Membranes were imaged on an iBright FL1500 (590 nm emission), and total protein (No-Stain Labeling Buffer, Invitrogen) was used as the normalisation control.

### Targeted insulin nanoparticle treatment

Glucose-responsive nanoparticles enable insulin delivery proportional to circulating glucose levels, allowing assessment under dynamic metabolic conditions. Glucose-responsive nanoparticles (PG_EDA_-_FPBA_) were synthesised as reported previously(*46*). Briefly, Phytoglycogen (PG) (200 mg) was dissolved in acetic buffer (5 mL, 0.6 M, pH 5.5) and mixed with sodium periodate (42 mg, 0.24 mmol) in the dark for 2 h. EDA (72 mg, 1.2 mmol) and sodium cyanoborohydride (12 mmol) were added and stirred overnight. The product (PG_EDA_ NPs) was purified by dialysis (molecular weight cutoff (MWCO) 14 kDa) against Milli-Q water for 3 days and then freeze-dried. FPBA (40 mL; 110 mg, 0.6 mmol), *N*-hydroxysuccinimide (NHS; 140 mg, 1.2 mmol) and *N*-(3-dimethylaminopropyl)-*N*′-ethylcarbodiimide hydrochloride (EDC•HCl; 120 mg, 0.6 mmol) were combined over 30 min. PG_EDA_ NPs (100 mg) were added to the reaction mixture and stirred overnight in dark. The final product (PG_EDA-FPBA_ NPs) was purified by dialysis (MWCO cutoff 14 kDa) against NaCl (0.1 M) for 1 day and Milli-Q water for 3 days and freeze-dried. The PG_EDA_-_FPBA_ NPs were characterised by ^1^H-NMR spectroscopy, absorption spectroscopy, and DLS. Characterisation of glucose binding capacity and insulin release was previously reported(*46*).

Mice (WT and *mdx*) were fasted for ∼5 h, and body weight was measured. Those mice were then subcutaneously injected with glucose-responsive nanoparticles (control) or glucose-responsive nanoparticle loaded with insulin (insulin dose, 22.5 IU/kg). Baseline glucose tests and glucose tolerance tests were conducted as described above at intervals 1 day, 1 week and 1 month post injection of nanoparticles. Mouse tissues (brain, heart, lung, liver, kidney, spleen, and skeletal muscle) were collected following euthanasia by cervical dislocation. Organs were rinsed in cold DPBS and fixed in 10% neutral-buffered formalin overnight. Fixed tissues were paraffin-embedded, sectioned at 4 µm, mounted on Superfrost slides, and stained with haematoxylin and eosin. Stained sections were imaged using an Olympus BX51 fluorescence microscope(*57*).

The mice were euthanised by cervical dislocation, and major organs (including the brain, heart, lung, liver, kidney, spleen, and muscle) were harvested. Mouse tissues were collected and rinsed in cold DPBS, then fixed in 10% formalin overnight. Tissues were paraffin-embedded, sectioned at 4 microns, and floated sections into superfrost slides and then subjected to H&E staining. An Olympus BX51 fluorescence microscope was used to image the H&E-stained tissue slides [PMID: 41165268].

### Promethion metabolic system studies

Mice were singularly housed in the Promethion Metabolic System for synchronous measurement of VO_2_, VCO_2_, energy expenditure and activity levels over 1h. The assessment included an initial 1h acclimation period before the data recording period. Treatment mice received an insulin (0.50 IU insulin/g body mass) immediately prior to a 30 s scruff-restraint stressor and then were placed into the Promethion system. A brief (30 s) scruff restraint was used to capture rapid physiological and behavioural responses, consistent with established paradigms in *mdx* mice(*33*, *34*, *47*). Control mice were not injected with insulin but were scruff-restrained and placed into the Promethion system. All mice were able to freely ambulate around the cage for the data recording period. Following completion of the experiment, mice were returned to their home cages.

### EMPA study – treatment regimen

To validate the effect of EMPA on blood glucose levels, 8-week-old *mdx* mice were treated with 300 mg/kg EMPA in chow for one week. After this time, mice were fasted and an intraperitoneal glucose tolerance test (IGTT) was conducted. Blood glucose was measured at 0, 15, 30, 45, 60, 90 and 120 min post-glucose administration (2 mg/g) using a LifeSmart glucometer (Model LS-946N, Genesis Biotech, Queensland, Australia). Vehicle-fed WT and *mdx* mice were used a genotype and treatment controls. The therapeutic potential of EMPA was then assessed short-term. At 3 weeks of age, WT mice were given a standard control diet (vehicle), whereas *mdx* mice were randomly allocated into: (1) standard chow (vehicle); (2) chow with EMPA (300 mg/kg); (3) chow with prednisolone (30mg/kg); or (4) chow with EMPA and prednisolone (all diets provided by Boehringer Ingelheim (Ingelheim, Germany) and manufactured by Ssniff Spezialdiäten GmbH). The dosage of EMPA was based on murine studies showing significant skeletal muscle effects in a model of heart failure(*58*).

### EMPA study – muscle characterisation

After 3 weeks of treatment, two-paw grip strength and whole-body strength were measured as previously described(*59*, *60*) and mice were fasted to quantify blood glucose and ketone levels using the LifeSmart glucometer. Mice were then anesthetised and urine was collected directly from the bladder to measure urinary glucose levels. The soleus was excised tendon to tendon and *ex vivo* muscle contractile properties were determined as described previously(*59*, *60*). The contralateral soleus was embedded in OCT, cryosectioned at 10µm, stained with haematoxylin and eosin and analysed using ImageJ as previously described(*59*, *60*).

### EMPA study – proteomics

The gastrocnemius was immediately snap frozen and TMT labelling proteomics were performed by Monash Proteomics and Metabolomics Platform as previously described(*61–63*). Briefly, frozen muscle was cryo-pulverised, and proteins were extracted, reduced, alkylated and digested using the S-Trap workflow according to the manufacturer’s instructions (Protifi)(*64*). Peptides were labelled using TMT 18-plex reagents according to the manufacturer’s protocol with channel isotope purity considered for placement of samples based on condition (Thermo Fisher Scientific; lot XJ351218 for 16 channels and lot YB368953 for two channels). The 126 channel was reserved as a pooled internal reference consisting of an equal mixture of all samples across the three plexes. Following labelling, samples were fractionated and pooled into 12 fractions for LC–MS/MS analysis. Digestion efficiency exceeded 95%, and TMT labelling efficiency exceeded 97% across all three plexes.

Peptide fractions were analysed on an Orbitrap Eclipse Tribrid mass spectrometer coupled to a Vanquish Neo nanoLC system (Thermo Fisher Scientific). The acquisition method was modified from previous analyses by incorporating a five-peptide close-out strategy for identification and applying a 1% false discovery rate (FDR) threshold per fraction (Comet search engine with xCorr of 1.4). Raw files were processed in Proteome Discoverer v3.1 (Thermo Fisher Scientific) using SEQUEST HT, Comet and MS Amanda 3.0 search engines, with INFERYS-based rescoring, against the reviewed *Mus musculus* Swiss-Prot proteome accessed in June 2023 and supplemented with common contaminants. Identification was controlled at 1% FDR at the peptide-spectrum match, peptide and protein levels. Protein quantification was performed using TMT reporter ion abundances, with lot-specific reporter ion isotope impurity corrections applied using the manufacturer-provided correction factors.

Quantitative statistical analysis was performed using TMT-Analyst within the Monash Analyst Suites(*65–67*). Standard processing parameters were applied, including internal reference scaling based on the pooled 126 channel to correct inter-plex batch effects. Significant proteins identified from this workflow were then used for downstream pathway analyses in ShinyGO (v0.81)(*68*) to reveal KEGG pathway enrichment of significant proteins (as previously described(*62*, *63*)), and Reactome (v94) for pathway analysis. The FDR of <0.05 is reported except for the GO pathways of interest where significance is set as p<0.05.

### Statistics

Full statistical outputs and raw data are provided in the Supplementary Data file (Excel format). Data were analysed using a combination of parametric and non-parametric approaches tailored to the experimental design. Longitudinal measurements - including GTT, ITT, and acute repeated stress paradigm - were primarily assessed using repeated measures ANOVA or Linear Mixed-Effects Models (LMM) with Bonferroni post-hoc adjustments. Crossover designs were employed for experiments comparing basal glucose, GTT, and FDG-PET imaging under stressed versus non-stressed conditions, and insulin versus no insulin conditions, to account for inter-subject variability. For FDG-PET imaging, left and right brain regions were averaged prior to LMM analysis. For multi-factor comparisons, such as between genotype or treatment, two-way ANOVA was utilised with Bonferroni post-hoc adjustments. Areas under the curve (AUC) were calculated via the trapezoidal rule; total and incremental AUCs were compared using independent samples t-tests, while population-level AUC differences were determined via permutation tests (1,000 iterations). EMPA proteomics data was quantified as detailed in(*62*). In cases where normality or variance assumptions were violated (e.g., EMPA parameters, islet area), Kruskal-Wallis with Dunn’s post-hoc, Welch’s ANOVA, or log-transformations were applied.

## Acknowledgments

We thank the Virginia Tech metabolism core for providing instrumentation and technical support. We thank the Vanderbilt University’s Mouse Metabolic Phenotyping Centre for *in vivo* double cannulation experiments. We thank the Monash Metabolic Phenotyping Platform for *in vivo* metabolic experiments. The authors acknowledge the facilities, and scientific and technical assistance of the National Imaging Facility (NIF), a National Collaborative Research Infrastructure Strategy (NCRIS) capability at Monash Biomedical Imaging (MBI), a Technology Research for bio-imaging experiments. This study used BPA-enabled (Bioplatforms Australia) / NCRIS-enabled (National Collaborative Research Infrastructure Strategy) infrastructure located at the Monash Proteomics and Metabolomics Platform. This work was further supported by Monash eResearch capabilities, including Research Data Storage and Nectar Research Cloud.

## Data, code, and materials availability

All data and statistical information used in this study are provided within the article and as a source data file. The proteomics dataset generated in this manuscript has been deposited on the PRIDE repository.

## Author contributions

Conceptualisation: A.L, G.S.M, C.E.H, F.C, K.C, A.P.R, C.A.G, A.H, E.R, C.P.N

Investigation: A.L, G.S.M, C.A.T, H.L, J.C, L.B, N.G, S.K, E.vdB, E.S, J,E, G.Z, R.X, S.K.B, T.V.H.M, J.R.S, R.B.S, D.L.M, L.L, M.dV

Visualisation: G.S.M, C.A.T, H.L, E.vdB, M.dB,

Writing—original draft: G.S.M

Writing—review & editing: G.S.M, A.L, C.E.H, F.C, K.C, A.P.R, C.A.G, A.H, E.R, C.P.N, C.A.T, H.L, J.C, L.B, N.G, S.K, E.vdB, E.S, J,E, G.Z, R.X, S.K.B, T.V.H.M, J.R.S, R.B.S, D.L.M, L.L, M.dV

