## Supplemental figures and tables for "Stress unmasks impaired endocrine coordination of glucose metabolism in dystrophin deficiency"

**SUPPLEMENTARY DATA**

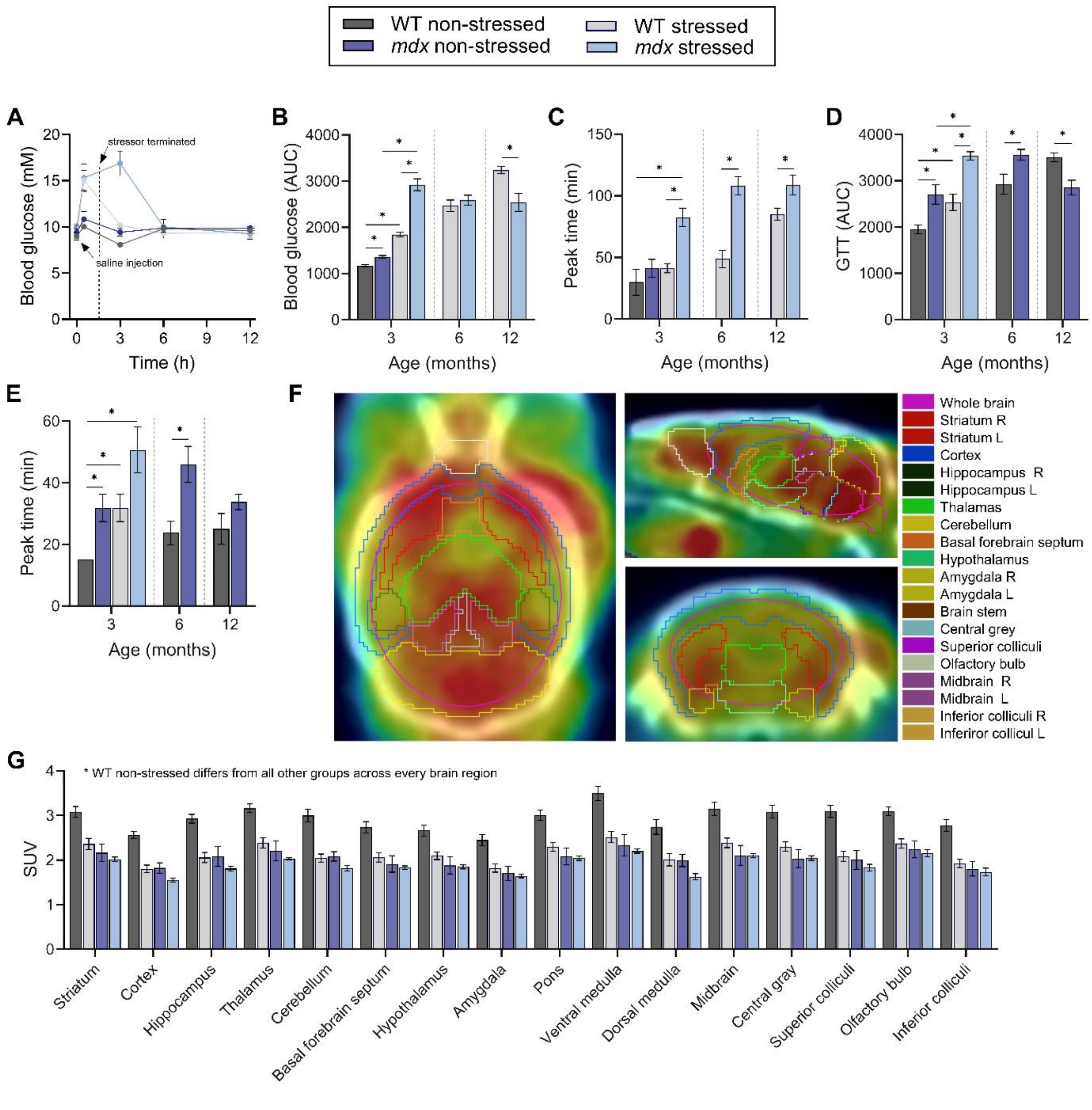

**Fig. S1.**

**Systemic glucose handling in *mdx* and wildtype (WT) mice. a,** Blood glucose concentrations measured over 12 h in 3-month-old, unfasted animals following an intraperitoneal injection of 0.9% saline at rest (non-stressed) or during a 2 h tube-restraint (stressed) with continued measurements 1, 4 and 10 h after stress termination. **b,c,** Area under the curve (AUC) **(b)** and time to peak blood glucose concentration **(c)** at 3, 6 and 12 months of age following a 5-h fast and intraperitoneal injection of 0.9% saline (procedural control). **d,e,** Area under the curve (AUC) **(d)** and time to peak blood glucose **(e)** at 3, 6 and 12 months of age following a 5-h fast and intraperitoneal injection of 2 mg $g^{-1}$ glucose (glucose tolerance test (GTT)). **f,** Representative [$^{18}$F] fluorodeoxyglucose ($^{18}$F-FDG) positron emission tomography (PET) scan matched to the Ma_Benveniste_Mirrione brain template with volumes of interest illustrated (coronal, sagittal and horizontal planes). **g,** $^{18}$FDG standardised uptake values (SUV) in templated brain regions. Data were analysed using repeated-measures ANOVA or linear mixed-effects models with Bonferroni post-hoc adjustments, or One-way ANOVA, Wilcoxon rank sum test or Kruskal-Wallis with Dunn's post-hoc testing. n = 5-8 per group. Data are mean ± SEM. * $P < 0.05$. Full statistical analyses and underlying raw data are provided in the Supplementary Data file.

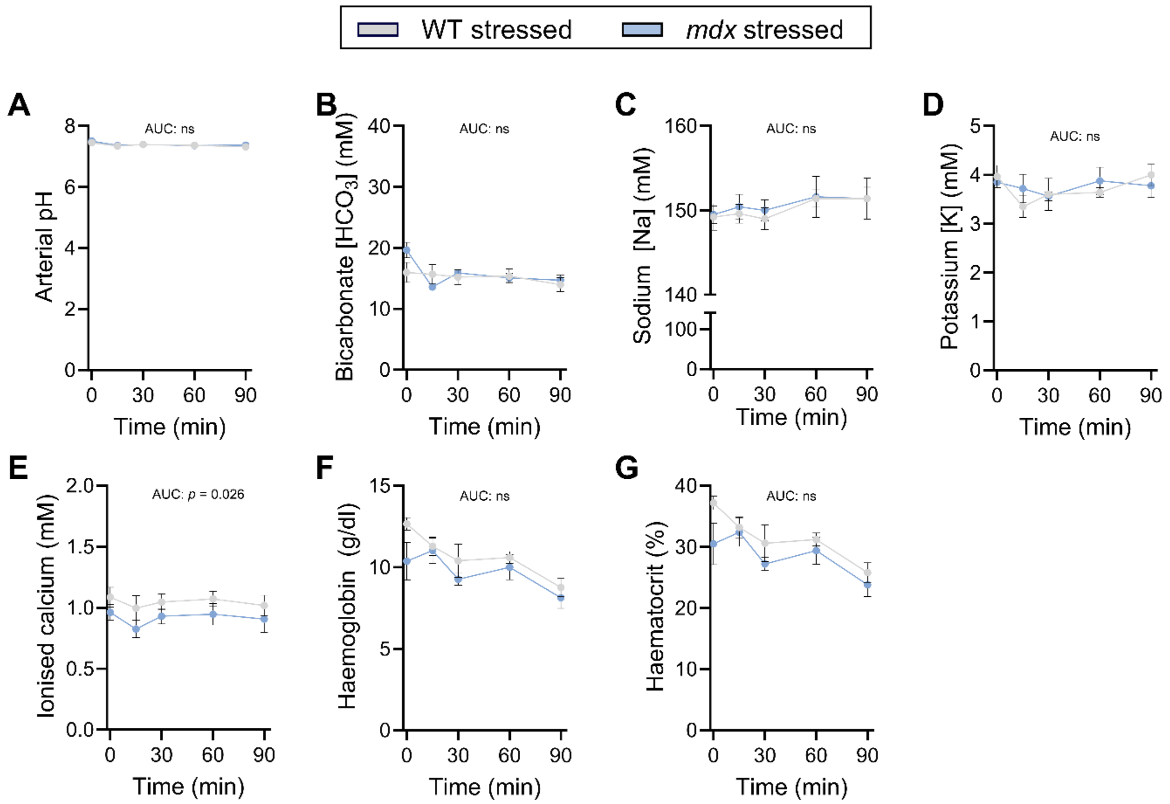

**Fig. S2.**

**Arterial blood chemistry during stress in *mdx* and wildtype (WT) mice. A-G,** Arterial blood pH **(A)**, bicarbonate **(B)**, sodium (Na) **(C)**, potassium (K) **(D)**, ionised calcium (Ca) concentration **(E)**, haemoglobin concentration **(F)**, and haematocrit **(G)** measured during a 90-minute tube-restraint stress. Data were analysed using linear mixed-effects models with Bonferroni post-hoc adjustments. n =5-10 per group. Data are mean ± SEM. $P < 0.05$. Full statistical analyses and underlying raw data are provided in the Supplementary Data file.

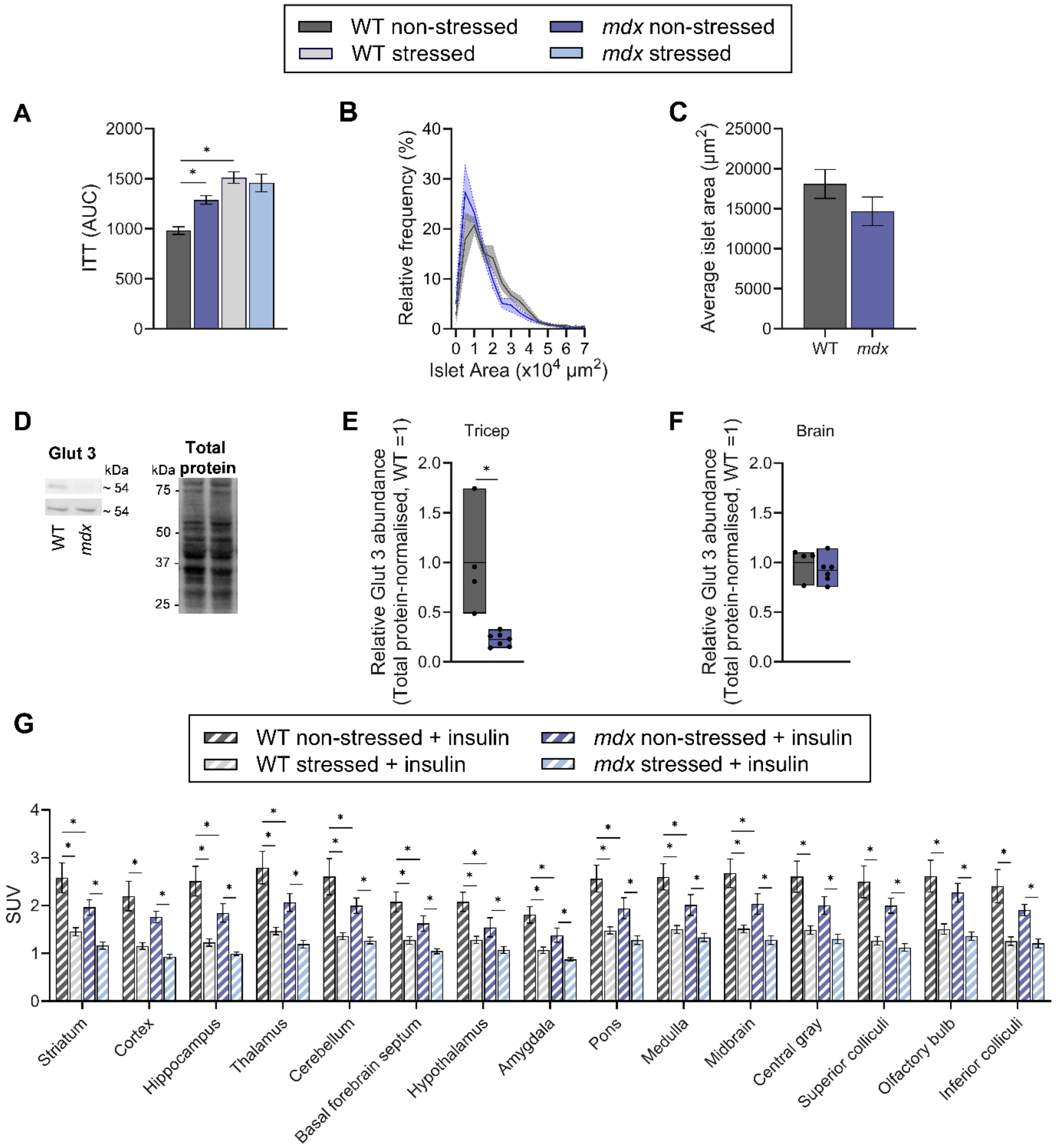

**Fig. S3.**

**Insulin-glucose crosstalk in *mdx* and wildtype (WT) mice. A,** Area under the curve (AUC) for blood glucose concentrations following a 5-h fast and intraperitoneal injection of insulin (0.5 IU kg$^{-1}$; insulin tolerance test (ITT)) measured at rest (non-stressed) or during a 2-h tube-restraint (stressed). **B,** Relative frequency distribution of pancreatic islet area from isolated islets. **C,** Mean pancreatic islet area. **D-F,** Representative immunoblot **(D)** and quantification of glucose transporter 3 (GLUT3) protein abundance in triceps **(E)** and brain **(F)** extracts, normalised to total protein and expressed relative to WT; bars indicate mean with minimum–maximum values. **G,** [$^{18}$F] fluorodeoxyglucose ($^{18}$F-FDG) standardised uptake values (SUV) in templated brain regions from mice treated with exogeneous insulin prior to stress or non-stress conditions. Data were analysed using repeated-measures ANOVA or linear mixed-effects models with Bonferroni post-hoc adjustments, or One-way ANOVA or Kruskal-Wallis with Dunn's post-hoc testing. n =4-8 per group. Data are mean ± SEM (unless otherwise stated). * $P < 0.05$. Full statistical analyses and underlying raw data are provided in the Supplementary Data file.

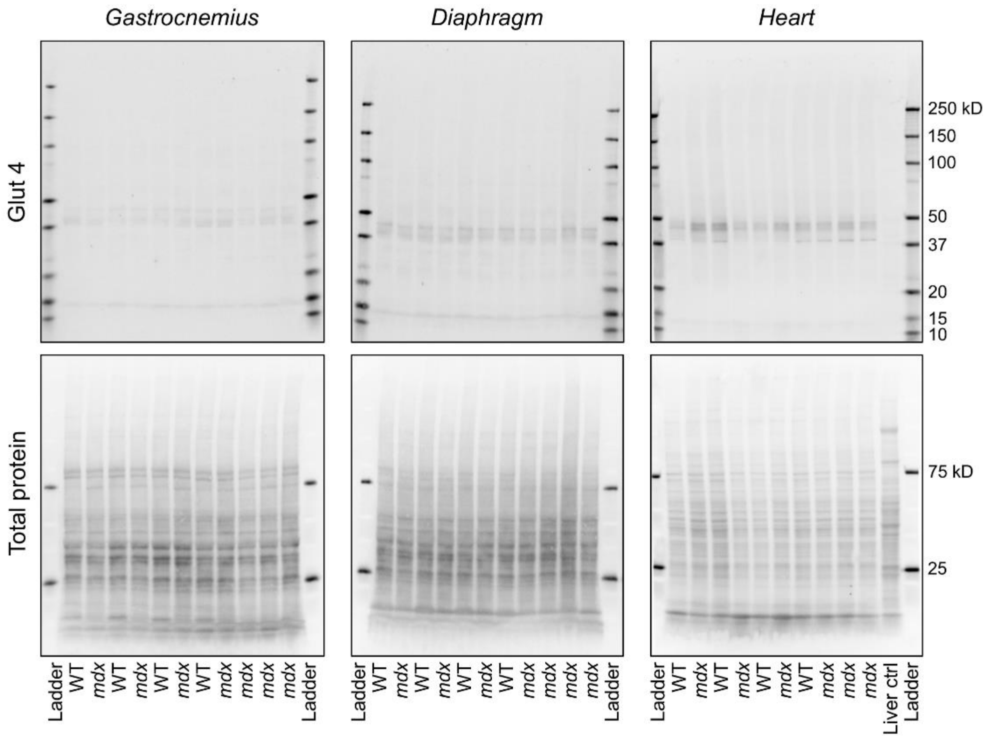

**Fig. S4.**

**Full-length immunoblots of glucose transporter 4 (Glut4) with total protein loading across gastrocnemius, diaphragm and heart tissues.**

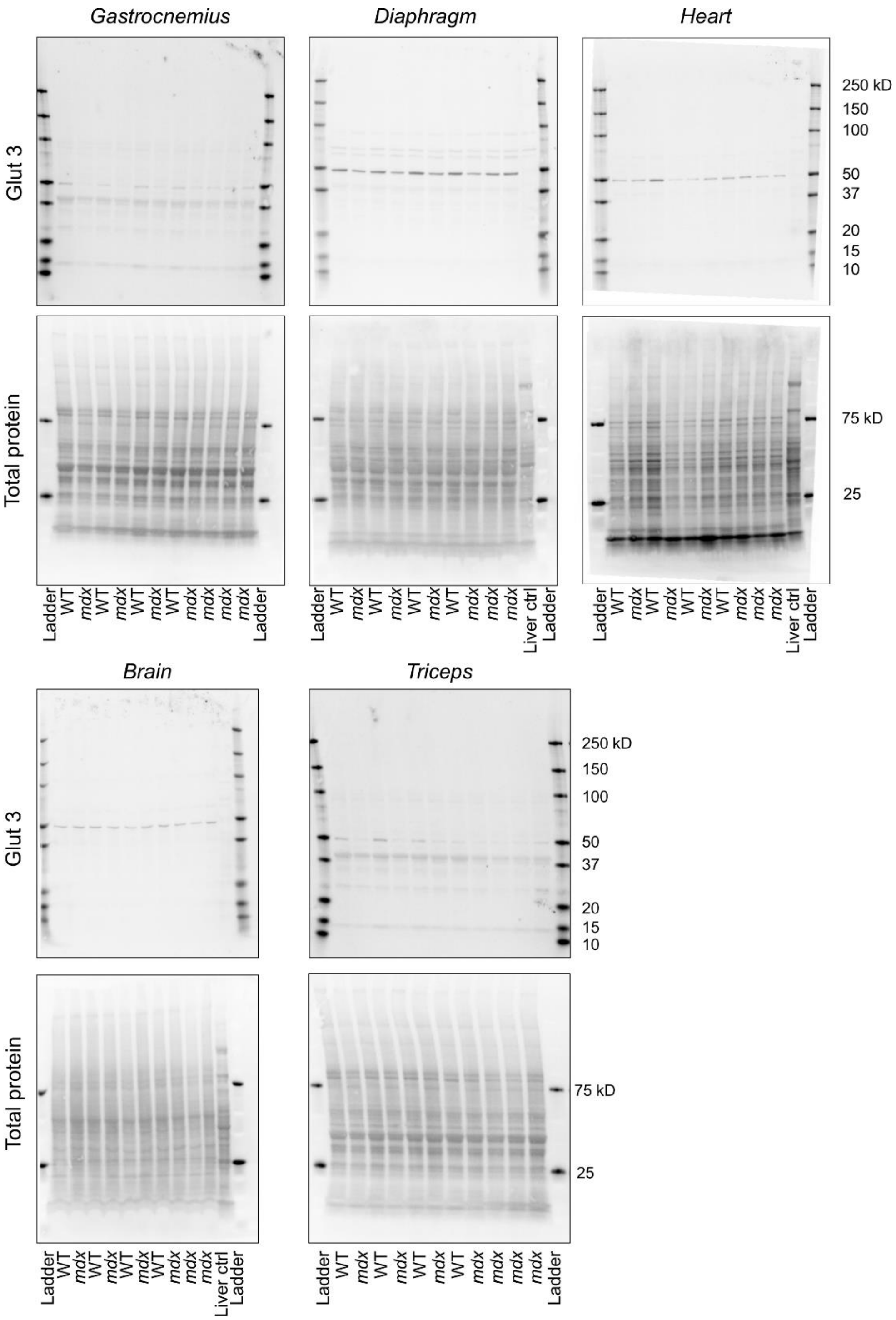

**Fig. S5.**

**Full-length immunoblots of glucose transporter 3 (Glut3) with total protein loading across gastrocnemius, diaphragm, heart, brain and triceps tissues.**

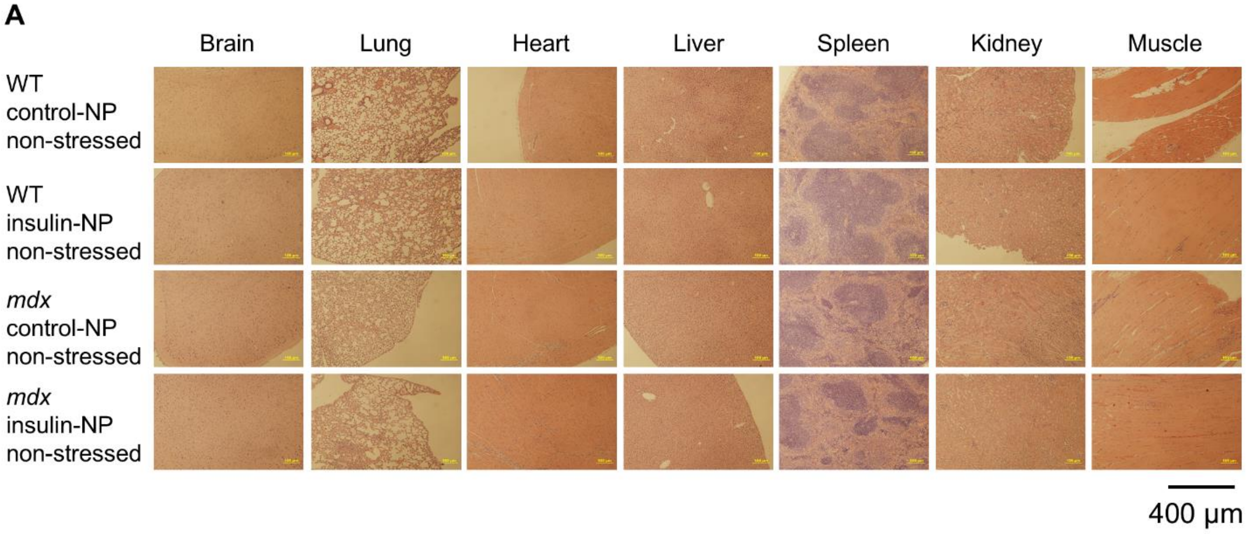

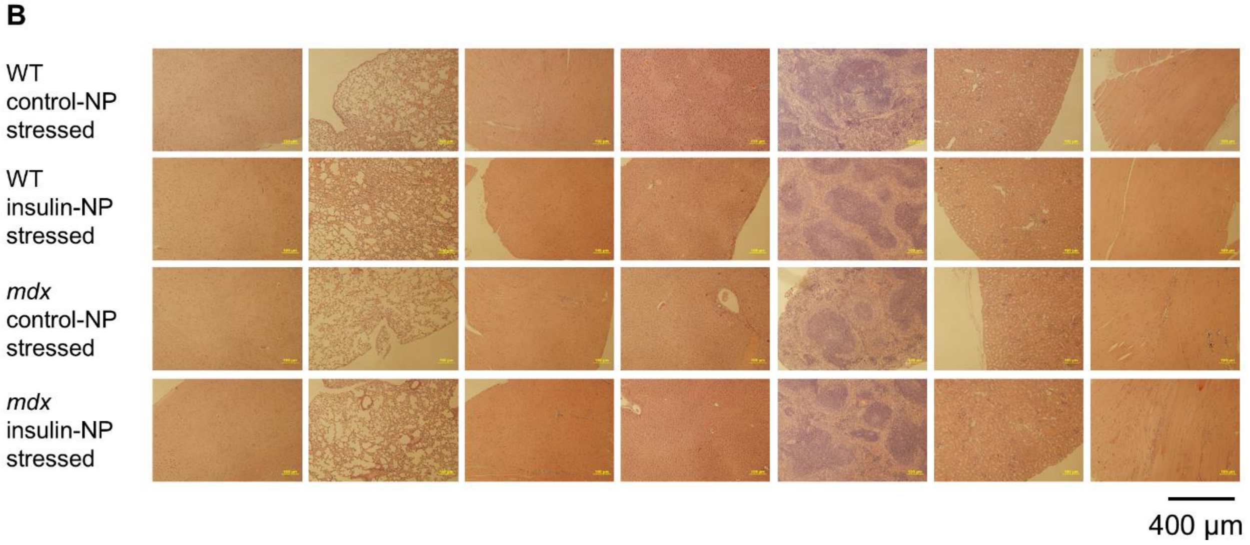

**Fig. S6.**

**The glucose-responsive insulin nanoparticle formulation showed a favourable safety profile**. **A,B,** Representative H&E histological samples of different tissues 1 month post subcutaneous injection of nanoparticles and basal glucose measures in non-stressed **(A)** or stressed **(B)** conditions**.**

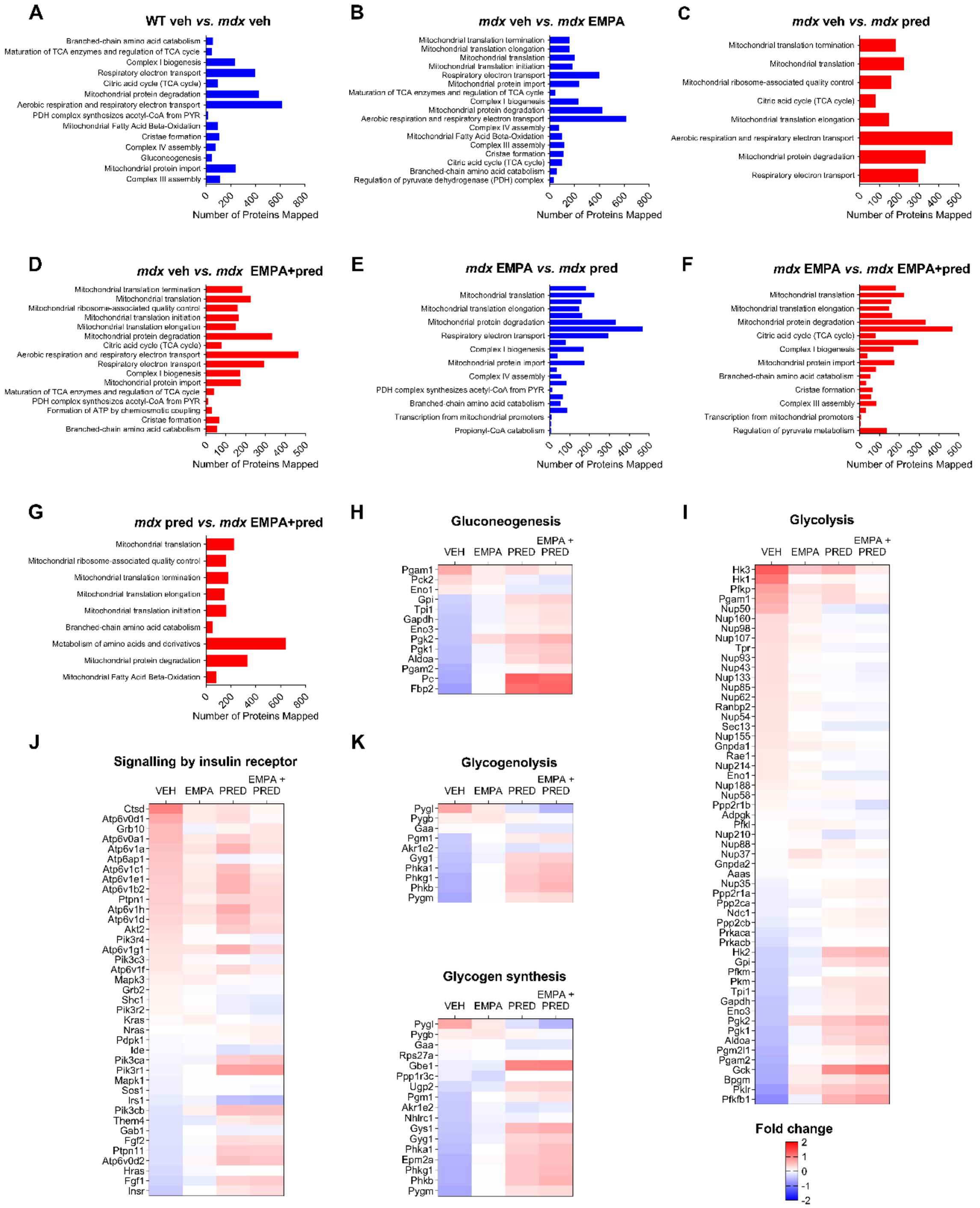

**Fig. S7.**

**Alterations to muscle metabolic pathways at the proteomic level. a-g,** Reactome pathway enrichment maps of metabolic pathways in skeletal muscle, showing the number of mapped proteins and direction of regulation for the indicated comparisons: **a,** WT vehicle (veh) vs. *mdx* veh; **b,** *mdx* veh vs. *mdx* empagliflozin (EMPA); **c,** *mdx* veh vs. *mdx* prednisolone (pred); **d,** *mdx* veh vs. *mdx* EMPA + pred; **e,** *mdx* EMPA vs. *mdx* pred; **f,** *mdx* EMPA vs. *mdx* EMPA

+ pred; **g,** *mdx* pred vs. *mdx* EMPA + pred. Bar colour denotes direction of change (red, upregulation; blue, downregulation). **h-l,** Differential expression of proteins contributing to key GO metabolic pathways across treatment groups: **h,** gluconeogenesis; **i,** glycolysis; **j,** insulin receptor signalling; **k,** glycogenolysis; **l,** glycogen synthesis. Colour indicates fold change relative to comparator group (red, increased; blue, decreased).

**Table S1.**

**Significant KEGG pathway enrichment in skeletal muscle across treatment cohorts**. Enrichment analysis was filtered to include only pathways directly related to primary metabolism. Data are presented for *mdx* mice treated with vehicle (VEH), prednisolone (PRED) and a combination of Empagliflozin and Prednisolone (EMPA+PRED). Comparisons are defined as follow: the *mdx* VEH group is expressed relative to WT VEH and all other experimental groups are expressed relative to the *mdx* VEH group. No significantly enriched pathways were identified in the EMPA treatment group based on the established significance threshold (FDR < 0.05).

| **KEGG Pathway** | **Number of Hits** | **Fold Enrichment** | **-Log$_{10}$ of FDR** | **FDR** |
|---|---|---|---|---|
| *mdx* VEH | | | | |
| Metabolic pathways | 626 | 1.155 | 9.645 | <0.0001 |
| Oxidative phosphorylation | 93 | 1.345 | 6.109 | <0.0001 |
| Carbon metabolism | 79 | 1.374 | 6.109 | <0.0001 |
| Biosynthesis of amino acids | 45 | 1.399 | 3.685 | <0.001 |
| Citrate cycle (TCA cycle) | 23 | 1.461 | 2.349 | 0.004 |
| Fatty acid degradation | 31 | 1.372 | 1.977 | 0.011 |
| Propanoate metabolism | 26 | 1.407 | 1.977 | 0.011 |
| Glycolysis/Gluconeogenesis | 40 | 1.298 | 1.601 | 0.025 |
| Biosynthesis of nucleotide sugars | 27 | 1.361 | 1.510 | 0.031 |
| Amino sugar and nucleotide sugar metabolism | 35 | 1.311 | 1.506 | 0.031 |
| Pyruvate metabolism | 30 | 1.328 | 1.396 | 0.040 |
| *mdx* PRED | | | | |
| PPAR signalling pathway | 32 | 1.578 | 1.705 | 0.019 |
| *mdx* EMPA+PRED | | | | |
| Oxidative phosphorylation | 71 | 1.382 | 2.477 | 0.003 |
| Carbon metabolism | 59 | 1.381 | 2.032 | 0.009 |
| Fatty acid degradation | 26 | 1.548 | 1.694 | 0.020 |
| PPAR signalling pathway | 33 | 1.474 | 1.694 | 0.020 |

**Table S2.**

**Differential expression of proteins involved in the Reactome gluconeogenesis and glycogen synthesis pathways across treatment groups.** Data are presented for *mdx* mice treated with vehicle (VEH), prednisolone (PRED) and a combination of Empagliflozin and Prednisolone (EMPA+PRED). For each protein identified in the pathway, fold change (Log2FC), and the raw P-value are provided. Comparisons are defined as follow: the *mdx* VEH group is expressed relative to WT VEH and all other experimental groups are expressed relative to the *mdx* VEH group.

| | ***mdx* VEH** | | ***mdx* EMPA** | | ***mdx* PRED** | | ***mdx* EMPA+PRED** | |
|---|---|---|---|---|---|---|---|---|
| | **Log2FC** | ***P* Value** | **Log2FC** | ***P* Value** | **Log2FC** | ***P* Value** | **Log2FC** | ***P* Value** |
| *Gluconeogenesis* | | | | | | | | |
| **Pgam1** | 0.667 | <0.0001 | 0.156 | 0.064 | 0.295 | <0.001 | 0.123 | 0.140 |
| **Pck2** | 0.317 | <0.0001 | 0.150 | 0.055 | -0.117 | 0.131 | -0.224 | 0.005 |
| **Eno1** | 0.171 | <0.0001 | 0.042 | 0.297 | -0.167 | <0.001 | -0.173 | <0.001 |
| **Gpi** | -0.374 | <0.0001 | -0.098 | 0.182 | 0.288 | <0.001 | 0.356 | <0.0001 |
| **Tpi1** | -0.409 | <0.0001 | -0.102 | 0.183 | 0.175 | 0.025 | 0.238 | 0.003 |
| **Gapdh** | -0.448 | <0.0001 | -0.105 | 0.183 | 0.150 | 0.059 | 0.239 | 0.004 |
| **Eno3** | -0.471 | <0.0001 | -0.055 | 0.501 | 0.107 | 0.190 | 0.178 | 0.032 |
| **Pgk2** | -0.471 | <0.001 | 0.286 | 0.034 | 0.460 | 0.001 | 0.583 | <0.001 |
| **Pgk1** | -0.512 | <0.0001 | -0.086 | 0.264 | 0.364 | <0.0001 | 0.414 | <0.0001 |
| **Aldoa** | -0.532 | <0.0001 | -0.084 | 0.267 | 0.292 | <0.001 | 0.444 | <0.0001 |
| **Pgam2** | -0.629 | <0.0001 | -0.035 | 0.664 | 0.049 | 0.542 | 0.150 | 0.066 |
| **Pc** | -0.674 | <0.0001 | 0.009 | 0.926 | 1.250 | <0.0001 | 1.160 | <0.0001 |
| **Fbp2** | -0.770 | <0.0001 | 0.019 | 0.869 | 1.110 | <0.0001 | 1.210 | <0.0001 |
| *Glycogen synthesis* | | | | | | | | |
| **Pygl** | 0.687 | <0.0001 | 0.177 | 0.116 | -0.240 | 0.035 | -0.578 | <0.0001 |
| **Pygb** | 0.137 | 0.027 | 0.188 | 0.003 | 0.071 | 0.239 | -0.055 | 0.363 |
| **Gaa** | 0.038 | 0.438 | 0.008 | 0.875 | -0.206 | <0.001 | -0.200 | <0.001 |
| **Rps27a** | -0.056 | 0.115 | -0.011 | 0.750 | -0.042 | 0.234 | 0.015 | 0.662 |
| **Gbe1** | -0.137 | 0.101 | -0.156 | 0.061 | 0.950 | <0.0001 | 0.996 | <0.0001 |
| **Ppp1r3c** | -0.149 | 0.435 | -0.315 | 0.103 | 2.160 | <0.0001 | 2.400 | <0.0001 |
| **Ugp2** | -0.300 | <0.0001 | -0.187 | 0.003 | 0.329 | <0.0001 | 0.350 | <0.0001 |
| **Pgm1** | -0.411 | <0.0001 | -0.006 | 0.925 | 0.140 | 0.043 | 0.232 | 0.001 |
| **Akr1e2** | -0.426 | <0.0001 | -0.077 | 0.181 | -0.247 | <0.0001 | -0.178 | 0.003 |
| **Nhlrc1** | -0.436 | <0.0001 | -0.090 | 0.266 | -0.131 | 0.106 | -0.038 | 0.634 |
| **Gys1** | -0.454 | <0.0001 | -0.136 | 0.051 | 0.531 | <0.0001 | 0.640 | <0.0001 |
| **Gyg1** | -0.492 | <0.0001 | -0.111 | 0.117 | 0.327 | <0.0001 | 0.392 | <0.0001 |
| **Phka1** | -0.547 | <0.0001 | -0.005 | 0.954 | 0.382 | <0.0001 | 0.495 | <0.0001 |
| **Epm2a** | -0.611 | <0.0001 | 0.033 | 0.702 | 0.434 | <0.0001 | 0.488 | <0.0001 |
| **Phkg1** | -0.617 | <0.0001 | 0.005 | 0.954 | 0.434 | <0.0001 | 0.529 | <0.0001 |
| **Phkb** | -0.624 | <0.0001 | 0.010 | 0.898 | 0.439 | <0.0001 | 0.566 | <0.0001 |
| **Pygm** | -0.685 | <0.0001 | -0.015 | 0.855 | 0.243 | 0.005 | 0.303 | |

**Table S3.**

**Differential expression of proteins involved in the Reactome glycolysis and glycogenolysis pathways across treatment groups.** Data are presented for *mdx* mice treated with vehicle (VEH), prednisolone (PRED) and a combination of Empagliflozin and Prednisolone (EMPA+PRED). For each protein identified in the pathway, fold change (Log2FC), and the raw P-value are provided. Comparisons are defined as follow: the *mdx* VEH group is expressed relative to WT VEH and all other experimental groups are expressed relative to the *mdx* VEH group.

| | ***mdx* VEH** | | ***mdx* EMPA** | | ***mdx* PRED** | | ***mdx* EMPA+PRED** | |
|---|---|---|---|---|---|---|---|---|
| | **Log2FC** | ***P* Value** | **Log2FC** | ***P* Value** | **Log2FC** | ***P* Value** | **Log2FC** | ***P* Value** |
| *Glycolysis* | | | | | | | | |
| **Hk3** | 1.290 | <0.0001 | 0.428 | 0.037 | 0.627 | 0.003 | 0.157 | 0.435 |
| **Hk1** | 1.090 | <0.0001 | 0.043 | 0.496 | 0.071 | 0.263 | -0.034 | 0.593 |
| **Pfkp** | 0.745 | <0.0001 | 0.213 | 0.035 | 0.307 | 0.003 | 0.040 | 0.684 |
| **Pgam1** | 0.667 | <0.0001 | 0.156 | 0.064 | 0.295 | 0.001 | 0.123 | 0.140 |
| **Nup50** | 0.566 | <0.0001 | 0.114 | 0.131 | -0.162 | 0.034 | -0.276 | <0.001 |
| **Nup160** | 0.290 | <0.0001 | 0.127 | 0.011 | -0.054 | 0.264 | -0.025 | 0.599 |
| **Nup98** | 0.270 | <0.0001 | 0.050 | 0.203 | -0.050 | 0.200 | -0.073 | 0.063 |
| **Nup107** | 0.267 | <0.0001 | 0.106 | 0.023 | 0.030 | 0.516 | -0.012 | 0.800 |
| **Tpr** | 0.255 | <0.0001 | 0.069 | 0.038 | -0.077 | 0.021 | -0.116 | <0.001 |
| **Nup93** | 0.253 | <0.0001 | 0.042 | 0.240 | 0.008 | 0.816 | -0.012 | 0.739 |
| **Nup43** | 0.236 | <0.001 | -0.025 | 0.674 | -0.026 | 0.667 | -0.119 | 0.052 |
| **Nup133** | 0.236 | <0.0001 | 0.054 | 0.181 | -0.042 | 0.300 | -0.090 | 0.028 |
| **Nup85** | 0.232 | <0.0001 | 0.022 | 0.607 | -0.046 | 0.289 | -0.048 | 0.267 |
| **Nup62** | 0.232 | <0.0001 | 0.071 | 0.059 | 0.011 | 0.761 | -0.028 | 0.449 |
| **Ranbp2** | 0.223 | <0.0001 | 0.059 | 0.059 | -0.091 | 0.005 | -0.084 | 0.008 |
| **Nup54** | 0.222 | <0.0001 | 0.018 | 0.657 | 0.027 | 0.498 | -0.050 | 0.206 |
| **Sec13** | 0.220 | <0.0001 | 0.005 | 0.901 | -0.152 | <0.001 | -0.155 | <0.001 |
| **Nup155** | 0.211 | <0.0001 | 0.073 | 0.070 | 0.012 | 0.756 | -0.005 | 0.900 |
| **Gnpda1** | 0.186 | <0.001 | 0.140 | 0.009 | 0.085 | 0.102 | -0.030 | 0.554 |
| **Rae1** | 0.177 | <0.001 | -0.021 | 0.677 | -0.078 | 0.116 | -0.067 | 0.177 |
| **Nup214** | 0.173 | <0.001 | 0.073 | 0.109 | 0.006 | 0.899 | -0.052 | 0.247 |
| **Eno1** | 0.171 | <0.001 | 0.042 | 0.297 | -0.167 | <0.001 | -0.173 | <0.0001 |
| **Nup188** | 0.126 | 0.044 | 0.068 | 0.268 | 0.050 | 0.415 | -0.043 | 0.483 |
| **Nup58** | 0.087 | 0.056 | 0.026 | 0.568 | 0.065 | 0.153 | 0.036 | 0.426 |
| **Ppp2r1b** | 0.066 | 0.477 | -0.069 | 0.459 | -0.086 | 0.354 | -0.274 | 0.005 |
| **Adpgk** | 0.024 | 0.563 | 0.032 | 0.448 | -0.039 | 0.351 | -0.079 | 0.060 |
| **Pfkl** | 0.011 | 0.839 | 0.097 | 0.074 | 0.077 | 0.156 | -0.038 | 0.483 |
| **Nup210** | 0.002 | 0.964 | 0.052 | 0.134 | -0.244 | <0.0001 | -0.126 | <0.001 |
| **Nup88** | -0.022 | 0.661 | 0.041 | 0.404 | 0.102 | 0.042 | 0.014 | 0.769 |
| **Nup37** | -0.030 | 0.759 | 0.257 | 0.011 | 0.068 | 0.490 | 0.102 | 0.300 |
| **Gnpda2** | -0.031 | 0.625 | 0.084 | 0.188 | 0.046 | 0.469 | -0.055 | 0.382 |
| **Aaas** | -0.045 | 0.243 | -0.001 | 0.971 | -0.005 | 0.905 | -0.013 | 0.744 |
| **Nup35** | -0.116 | 0.035 | -0.008 | 0.880 | 0.065 | 0.232 | 0.120 | 0.029 |
| **Ppp2r1a** | -0.143 | 0.001 | -0.064 | 0.129 | 0.085 | 0.044 | 0.123 | 0.004 |
| **Ppp2ca** | -0.148 | 0.003 | -0.122 | 0.013 | -0.006 | 0.898 | 0.056 | 0.246 |
| **Ndc1** | -0.148 | 0.005 | 0.023 | 0.641 | 0.077 | 0.131 | 0.134 | 0.009 |
| **Ppp2cb** | -0.170 | <0.0001 | -0.063 | 0.065 | 0.066 | 0.053 | 0.109 | 0.002 |
| **Prkaca** | -0.244 | <0.001 | -0.147 | 0.024 | 0.008 | 0.904 | 0.039 | 0.538 |
| **Prkacb** | -0.284 | <0.0001 | -0.067 | 0.077 | -0.018 | 0.625 | -0.015 | 0.693 |
| **Hk2** | -0.371 | <0.0001 | -0.160 | 0.019 | 0.520 | <0.0001 | 0.568 | <0.0001 |

| | | | | | | | | |
|---|---|---|---|---|---|---|---|---|
| **Gpi** | -0.374 | <0.001 | -0.098 | 0.182 | 0.288 | <0.001 | 0.356 | <0.0001 |
| **Pfkm** | -0.386 | <0.0001 | -0.137 | 0.038 | -0.020 | 0.756 | 0.063 | 0.330 |
| **Pkm** | -0.406 | <0.0001 | -0.019 | 0.811 | 0.224 | 0.006 | 0.235 | 0.004 |
| **Tpi1** | -0.409 | <0.0001 | -0.102 | 0.183 | 0.175 | 0.025 | 0.238 | 0.003 |
| **Gapdh** | -0.448 | <0.0001 | -0.105 | 0.183 | 0.150 | 0.059 | 0.239 | 0.004 |
| **Eno3** | -0.471 | <0.0001 | -0.055 | 0.501 | 0.107 | 0.190 | 0.178 | 0.032 |
| **Pgk2** | -0.471 | <0.001 | 0.286 | 0.034 | 0.460 | 0.001 | 0.583 | <0.0001 |
| **Pgk1** | -0.512 | <0.0001 | -0.086 | 0.264 | 0.364 | <0.0001 | 0.414 | <0.0001 |
| **Aldoa** | -0.532 | <0.0001 | -0.084 | 0.267 | 0.292 | <0.001 | 0.444 | <0.0001 |
| **Pgm2l1** | -0.577 | <0.0001 | 0.034 | 0.564 | 0.089 | 0.140 | 0.208 | 0.001 |
| **Pgam2** | -0.629 | <0.0001 | -0.035 | 0.664 | 0.049 | 0.542 | 0.150 | 0.066 |
| **Gck** | -0.639 | <0.001 | 0.156 | 0.309 | 0.882 | <0.0001 | 1.060 | <0.0001 |
| **Bpgm** | -0.654 | <0.0001 | 0.134 | 0.349 | 0.223 | 0.121 | 0.346 | 0.018 |
| **Pklr** | -0.798 | <0.0001 | 0.259 | 0.149 | 0.444 | 0.015 | 0.493 | 0.007 |
| **Pfkfb1** | -0.910 | <0.0001 | -0.120 | 0.257 | 0.601 | <0.0001 | 0.732 | <0.0001 |
| *Glycogenolysis* | | | | | | | | |
| **Pygl** | 0.687 | <0.0001 | 0.177 | 0.116 | -0.240 | 0.035 | -0.578 | <0.0001 |
| **Pygb** | 0.137 | 0.027 | 0.188 | 0.003 | 0.071 | 0.239 | -0.055 | 0.363 |
| **Gaa** | 0.038 | 0.438 | 0.008 | 0.875 | -0.206 | <0.001 | -0.200 | <0.001 |
| **Pgm1** | -0.411 | <0.0001 | -0.006 | 0.925 | 0.140 | 0.043 | 0.232 | 0.001 |
| **Akr1e2** | -0.426 | <0.0001 | -0.077 | 0.181 | -0.247 | <0.0001 | -0.178 | 0.003 |
| **Phka1** | -0.547 | <0.0001 | -0.005 | 0.954 | 0.382 | <0.0001 | 0.495 | <0.0001 |
| **Phkg1** | -0.617 | <0.0001 | 0.005 | 0.954 | 0.434 | <0.0001 | 0.529 | <0.0001 |
| **Phkb** | -0.624 | <0.0001 | 0.010 | 0.898 | 0.439 | <0.0001 | 0.566 | <0.0001 |
| **Pygm** | -0.685 | <0.0001 | -0.015 | 0.855 | 0.243 | 0.005 | 0.303 | <0.001 |

**Table S4.**

**Differential expression of proteins involved in the Reactome insulin receptor signalling pathway across treatment groups.** Data are presented for *mdx* mice treated with vehicle (VEH), prednisolone (PRED) and a combination of Empagliflozin and Prednisolone (EMPA+PRED). For each protein identified in the pathway, fold change (Log2FC), and the raw P-value are provided. Comparisons are defined as follow: the *mdx* VEH group is expressed relative to WT VEH and all other experimental groups are expressed relative to the *mdx* VEH group.

| | ***mdx* VEH** | | ***mdx* EMPA** | | ***mdx* PRED** | | ***mdx* EMPA+PRED** | |
|---|---|---|---|---|---|---|---|---|
| | **Log2FC** | ***P* Value** | **Log2FC** | ***P* Value** | **Log2FC** | ***P* Value** | **Log2FC** | ***P* Value** |
| **Grb10** | 0.545 | <0.0001 | -0.117 | 0.159 | 0.083 | 0.318 | 0.191 | 0.024 |
| **Akt2** | 0.287 | <0.0001 | 0.039 | 0.367 | 0.404 | <0.0001 | 0.296 | <0.0001 |
| **Pik3r4** | 0.246 | <0.0001 | 0.028 | 0.590 | 0.006 | 0.903 | -0.117 | 0.029 |
| **Pik3c3** | 0.191 | 0.001 | 0.054 | 0.341 | -0.024 | 0.667 | -0.125 | 0.032 |
| **Mapk3** | 0.120 | 0.033 | 0.154 | 0.007 | 0.044 | 0.427 | -0.054 | 0.326 |
| **Grb2** | 0.092 | 0.005 | -0.025 | 0.424 | -0.079 | 0.015 | -0.070 | 0.031 |
| **Shc1** | 0.090 | 0.047 | -0.019 | 0.675 | -0.148 | 0.001 | -0.179 | <0.001 |
| **Pik3r2** | 0.071 | 0.423 | 0.033 | 0.705 | -0.147 | 0.098 | -0.199 | 0.028 |
| **Kras** | 0.033 | 0.499 | 0.103 | 0.041 | -0.002 | 0.961 | -0.082 | 0.097 |
| **Nras** | -0.007 | 0.783 | 0.005 | 0.850 | 0.065 | 0.017 | 0.109 | <0.001 |
| **Pdpk1** | -0.060 | 0.075 | -0.043 | 0.198 | 0.045 | 0.175 | 0.114 | 0.001 |
| **Pik3ca** | -0.086 | 0.158 | -0.091 | 0.136 | 0.390 | <0.0001 | 0.469 | <0.0001 |
| **Pik3r1** | -0.091 | 0.074 | 0.019 | 0.701 | 0.721 | <0.0001 | 0.751 | <0.0001 |
| **Mapk1** | -0.122 | <0.001 | -0.020 | 0.535 | 0.013 | 0.694 | 0.023 | 0.464 |
| **Sos1** | -0.130 | 0.012 | 0.012 | 0.812 | -0.022 | 0.662 | -0.064 | 0.206 |
| **Irs1** | -0.179 | 0.012 | -0.094 | 0.178 | -0.498 | <0.0001 | -0.539 | <0.0001 |
| **Pik3cb** | -0.232 | 0.001 | 0.113 | 0.097 | 0.514 | <0.0001 | 0.478 | <0.0001 |
| **Them4** | -0.238 | 0.006 | -0.196 | 0.022 | 0.158 | 0.063 | 0.237 | 0.006 |
| **Gab1** | -0.246 | <0.0001 | -0.007 | 0.876 | -0.119 | 0.015 | -0.108 | 0.026 |
| **Fgf2** | -0.256 | <0.0001 | -0.027 | 0.613 | 0.280 | <0.0001 | 0.252 | <0.0001 |
| **Ptpn11** | -0.264 | <0.0001 | -0.050 | 0.177 | 0.399 | <0.0001 | 0.392 | <0.0001 |
| **Atp6v0d2** | -0.272 | 0.079 | 0.049 | 0.748 | 0.478 | 0.003 | 0.437 | 0.006 |
| **Hras** | -0.305 | <0.0001 | -0.065 | 0.074 | 0.013 | 0.715 | 0.017 | 0.629 |
| **Fgf1** | -0.357 | <0.001 | -0.165 | 0.097 | 0.366 | <0.001 | 0.400 | <0.001 |
| **Insr** | -0.418 | <0.0001 | -0.010 | 0.836 | 0.192 | <0.001 | 0.281 | <0.0001 |
